# Turnover of RNA-binding Proteins and MicroRNAs by intrinsically disordered region-directed ZSWIM8 ubiquitin ligase during brain development

**DOI:** 10.1101/2024.01.27.577548

**Authors:** Jing Lei, Siming Zhong, Rong Fan, Xin Shu, Guan Wang, Jiansheng Guo, Shuting Xue, Luqian Zheng, Aiming Ren, Junfang Ji, Bing Yang, Shumin Duan, Zhiping Wang, Xing Guo

## Abstract

Widely present in mammalian proteomes, intrinsically disordered regions (IDRs) in proteins play important biological functions by conferring structural flexibility and mediating biomolecular interactions. IDR-containing proteins, including many RNA-binding proteins (RBPs), are prone to misfolding and aggregation and must be constantly monitored. Here we show that the conserved ZSWIM8-type Cullin-RING ubiquitin ligase (CRL^ZSWIM8^) is a master regulator of such proteins during brain development. ZSWIM8 selects its substrates via an IDR-dependent mechanism, and deletion of ZSWIM8 causes aberrant accumulation of numerous RBPs including AGO2 and ELAV1 in neonatal brains. Furthermore, AGO2 ubiquitination by ZSWIM8 is triggered by microRNA binding, leading to target-directed microRNA degradation (TDMD) of MiR7. Dysregulation of MiR7 in the absence of ZSWIM8 results in defects in oligodendrocyte maturation and functions. Together, our findings have demonstrated that, by utilizing variable target-recognition strategies, ZSWIM8 controls the abundance of conformationally flexible RBPs and miRNA metabolism that are essential for brain development.

**Teaser:** A conserved ubiquitin ligase controls the quality of disordered proteins to ensure brain development.

## Introduction

The structural information of a protein is not only manifested by folded globular domains but is also embedded in intrinsically disordered regions (IDRs). Biochemically, IDRs generally lack hydrophobic amino acids but possess an abundance of polar or charged residues, which reduces the propensity of forming a well-folded tertiary structure but confers structural flexibility. The vast majority of mammalian proteins are predicted to contain at least one recognizable IDR (*1*). The overall IDR content in mammalian proteomes has significantly increased through evolution, concomitant with the expansion of protein functions necessary for cell survival in higher organisms (*2–5*). IDRs play critical roles in mediating conformational changes and protein-protein interactions and are a common driving force of liquid-liquid phase separation (LLPS) (*1, 6, 7*). The same biochemical nature, however, also makes IDRs prone to folding errors and susceptible to aggregation, which are often hallmarks of neurodegenerative disorders (*8*). Moreover, when these proteins are mutated or misfolded, the deleterious defects can become contagious by perturbing functions of normal proteins that interact with them (*1*). Therefore, quality control of IDR-containing proteins is of utmost importance to cell survival and health.

A considerable portion of IDR-containing proteins are RNA-binding proteins (RBPs) that serve diverse and dynamic functions in shaping the cellular transcriptome (*9–11*). The number of repetitive tripeptides in IDRs of RBPs has expanded greatly from yeast to human, and such increase in RBP flexibility is expected to meet the ever-increasing complexity of RNA biology during evolution (*12*). In complex organs such as the brain, mRNA expression exhibits significant heterogeneity across different cell types, brain regions and developmental stages (*13, 14*). The spatiotemporal control of mRNA expression and function depends on RBPs as well as other RNAs (e.g., microRNAs), which themselves are tightly regulated by various miRNA-binding proteins (*9, 15–17*). IDR-containing RBPs such as TDP-43 and FUS are sensitive to stress and tend to form compact condensates in which their concentrations can become exceedingly high (*18*). Insoluble fibrillary tangles and amyloid plaques may then form, giving rise to the typical pathological features of neurodegenerative disorders such as amyotrophic lateral sclerosis (ALS) and frontotemporal lobar degeneration (FTLD) (*15, 19*). To maintain protein homeostasis in an IDR-abundant protein pool, eukaryotic cells have developed a complex protein quality control (PQC) system, including molecular chaperones and the ubiquitin-proteasome systems (UPS), to facilitate protein (re)folding and to clear detrimental proteins (*20, 21*). In the UPS pathway, proteins targeted for proteasomal degradation are often marked by polyubiquitination catalyzed by hundreds of E3 ubiquitin ligases.

Although a plethora of studies have established the substrate-enzyme relationship between numerous proteins and E3s (*22, 23*), how misfolding-prone IDR-containing proteins are specifically targeted remains poorly understood. Molecular chaperones such as Hsp90 can bind to a wide population of misfolded substrates and are believed to facilitate substrate recognition of specific E3 ligases (*20, 24*). Meanwhile, a limited number of E3 ligases are known to directly utilize their IDRs for recognition of IDR-enriched substrates. This so-called “disorder-targets-misorder” mechanism sets forth a unique principle underlying IDR recognition by E3s. In yeast, the E3 ubiquitin ligase San1 contains several conserved ordered motifs interspaced by large IDRs which provide an inherent plasticity for accommodating diverse misfolded substrates (*25*). Another IDR-containing yeast E3, CRL^Grr1^, was shown to degrade the IDR-containing protein Med13 in response to oxidative stress (*26*). In mammalian cells, the examples are limited to a few of E3s (e.g., CHIP and WWP2) and their individual substrates (IRF-1 and DVL-2) (*27, 28*). However, the generality of the “disorder-targets-misorder” mechanism of substrate recognition by mammalian E3s particularly *in vivo* has not been rigorously tested.

Previously we identified that EBAX-1 (Elongin B/C-Binding Axon Regulator), a novel substrate recognition subunit of Cullin-RING E3 ubiquitin ligases (CRL), guards neurodevelopment accuracy through quality control of the IDR-enriched axon guidance receptor SAX-3 in *C. elegans* (*24*). Later, we reported that the mammalian ortholog of EBAX-1, namely ZSWIM8, preferably targets the IDR region of DAB1, an adaptor downstream of REELIN, thereby safeguarding migration of neural progenitor cells in the developing mouse brain (*29*). Despite the N-terminal BC-box, CUL2-box and SWIM motif with known functions, the majority of ZSWIM8 C-terminal sequence is disordered, with three large IDRs punctuated with ordered sequences. In *C. elegans* and *Drosophila*, genetic mutations that create truncated proteins losing all or most of the C-terminal IDRs caused loss-of-function phenotypes, suggesting that these regions are indispensable for the EBAX-1/ZSWIM8 family (*24, 30*). These studies converge on the notion that ZSWIM8 and its homologs may use the “disorder-targets-misorder” mechanism for general substrate binding. Given that whole-body and brain-specific knockout of ZSWIM8 lead to developmental lethality in mice (*29, 31, 32*), we predict that ZSWIM8 may have a much broader substrate specificity in mammals.

Two recent studies uncovered a new role of ZSWIM8 in target-directed microRNA degradation (TDMD) of MiR7 in various cell lines (*33, 34*) through targeting Argonaute 2 (AGO2), a core component of the RNA-induced silencing complex (RISC) (*35–37*). In mice, MiR7 is encoded by *Mir7a-1*, *Mir7a-2* and *Mir7b* genes located on three different chromosomes (*38, 39*). As the primary source of mature MiR7, MiR7a shows broad expression in the cortical plate and ventricular zone of embryonic brains (*40*). Simultaneous suppression of MiR7 precursors causes dysregulated expression of critical transcription factors including TP53 and PAX6, and consequentially results in defective neurogenesis and microcephaly-like brain defects at birth (*40, 41*). Meanwhile, an *in vitro* study reported that intrinsically programmed downregulation of MiR7 in oligodendrocyte precursor cells (OPCs) is important for proper fate transition of OPCs into mature oligodendrocytes (OLs), which generate insulating myelin sheath surrounding neuronal axons to facilitate fast electrical conduction (*42*). These findings start to elucidate pivotal physiological roles of ZSWIM8 and MiR7 in the central nervous system (CNS). However, a few important questions remain unsolved. First, as a typical IDR-enriched E3 ligase, which population of substrates does ZSWIM8 target *in vivo*? Second, how does ZSWIM8 recognize disordered substrates (e.g., ROBO and DAB1) as well as structured substrates (e.g., AGO2)? Third, how does ZSWIM8-mediated TDMD of MiR7 participate in brain development?

Here, we provide comprehensive evidence demonstrating that ZSWIM8 is a versatile regulator of IDR-enriched proteins *in vivo*. Conditional deletion of ZSWIM8 in the developing mouse brain results in accumulation of miRNA-binding proteins (miRBPs) and upregulation of MiR7, accompanied by aberrant expression of proteins important for neuronal and glial functions. In particular, we have shown an unprecedented mechanism that MiR7 binding to AGO2 triggers ZSWIM8-mediated AGO2 ubiquitination. Additionally, we have demonstrated a new role of ZSWIM8-dependent TDMD in oligodendrocyte development. Deletion of ZSWIM8 causes accumulation of MiR7 in the oligodendrocyte lineage cells, which interferes the fate transition of OPCs and eventually impairs oligodendrocyte maturation and myelination. These findings have expanded our understanding about substrate recognition mechanisms of IDR-containing E3 ubiquitin ligases and uncovered the pivotal role of ZSWIM8-dependent TDMD in brain development.

## Results

### Deletion of ZSWIM8 in the nervous system causes broad accumulation of IDR-containing proteins

To define the spectrum of ZSWIM8 substrates in the mouse nervous system, we analyzed the whole proteomes of forebrain tissues dissected from *Zswim8^f/f^* and *Zswim8^f/f^;Nestin-Cre* (designated as *Z^f/f^* and *Z^f/f^;N-Cre* hereafter) littermates at birth (P0) or at postnatal day 14 (P14) (Fig. 1A and Fig. S1A).

**Figure 1.**
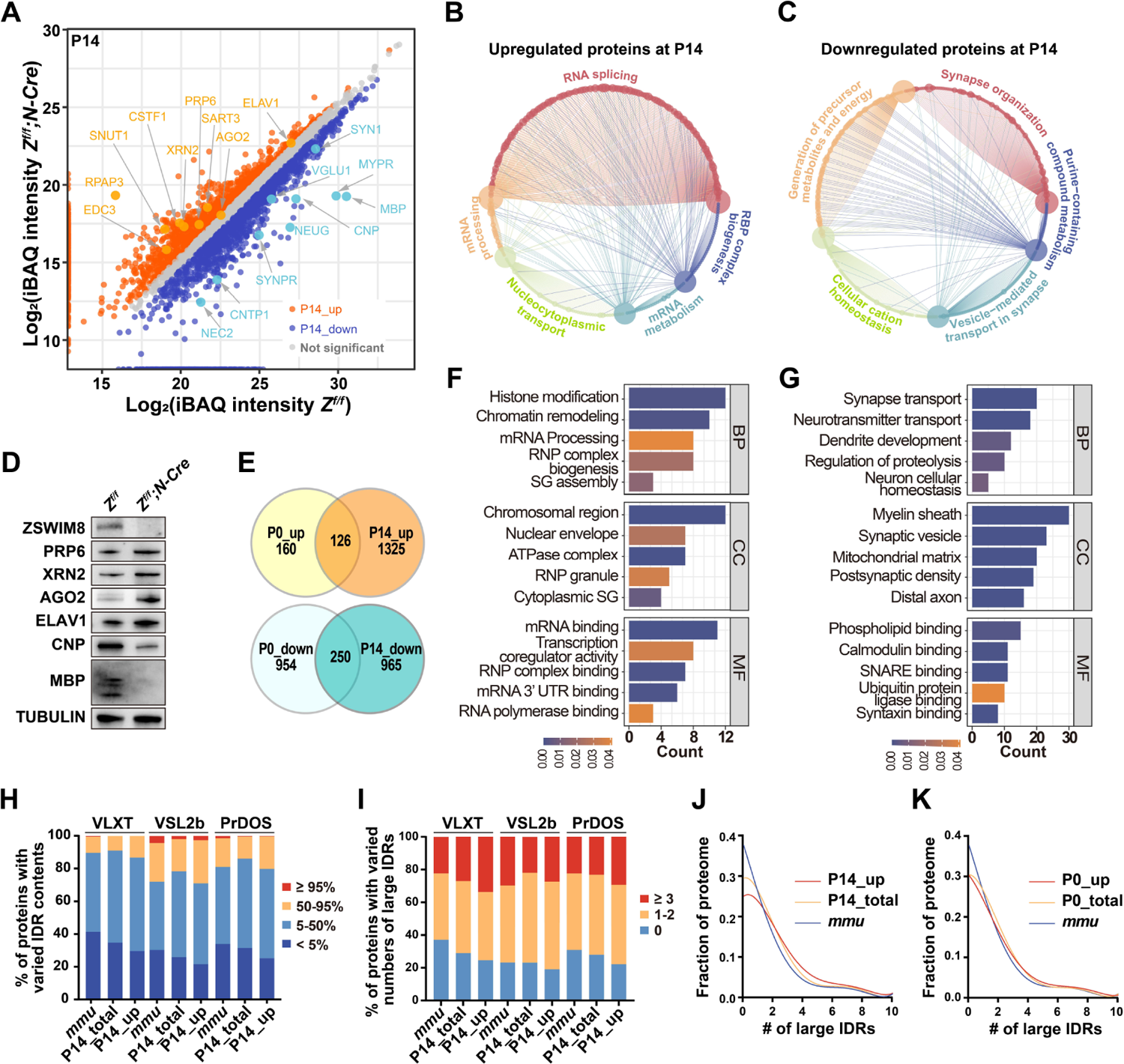
Deletion of ZSWIM8 in the mouse nervous system causes accumulation of proteins with IDRs. **(A)** P14 forebrain homogenates from *Z^f/f^* and *Z^f/f^;N-Cre* animals (N = 3 pairs of littermates) were analyzed by mass spectrometry. Representatives of differentially expressed proteins in *Z^f/f^;N-Cre* mice are indicated by arrows. Examples of RNA-binding proteins in the upregulated pool and proteins involved in neuronal and glial functions in the downregulated pool are highlighted by yellow dots and cyan dots, respectively. Mass spectrometry data of *Z^f/f^* and *Z^f/f^;N-Cre* samples were analyzed by the iBAQ (intensity-based absolute quantification) method and plotted. **(B, C)** Gene ontology (GO) analyses of upregulated (B) and downregulated proteins (C) in *Z^f/f^;N-Cre* animals at P14. Five enriched biological processes in each group are shown. **(D)** Western blot verification of the indicated proteins identified from the proteomic study of *Z^f/f^* and *Z^f/f^;N-Cre* samples at P14. PRP6, pre-mRNA processing factor 6; XRN2, 5’-3’ exoribonuclease 2; AGO2, argonaute RISC catalytic component 2; ELAV1, ELAV-like RNA binding protein; CNP, cyclic nucleotide phosphodiesterase; MBP, myelin basic protein. β-tubulin was used as loading control. **(E)** Venn diagrams comparing proteins up- or down-regulated at P0 and P14. **(F, G)** GO term analyses of proteins upregulated (F) or downregulated (G) in *Z^f/f^;N-Cre* animals at both P0 and P14 stages. BP, biological process; CC, cellular component; MF, molecular function. Color codes stand for Benjamini-Hochberg-adjusted p values. **(H)** Proteins in the total mouse proteome (*mmu*), in the forebrain proteome from control mice at P14 (P14_total), and those upregulated in *Z^f/f^;N-Cre* forebrain samples at P14 (P14_up) were ranked based on their IDR contents calculated by three algorithms (VXLT, VSL2b and PrDOS). **(I)** Proteins in the same groups as in (H) were ranked based on the number of large IDRs they harbor. **(J, K)** Fractional distributions of proteins with various numbers of large IDRs predicted by VLXT in the indicated proteomes. Data were fitted with the fifth order polynomial (R^2^ > 0.99).

We first focused on the P14 dataset, in which considerably more proteins were detected than from the P0 samples (4332 vs 2883 proteins). Gene ontology (GO) analyses indicated that in *Z^f/f^;N-Cre* brains, numerous proteins involved in RNA metabolism (e.g., ELAV1, AGO2, XRN2 and PRP6) were upregulated, whereas various proteins required for neuronal and glial functions (e.g., SYN1, NEUG, MBP and CNP) were downregulated (Fig. 1A-C and Fig. S1B and C). The differential expression of several proteins was further confirmed by western blot (Fig. 1D). In P0 forebrains, ZSWIM8 knockout also led to altered expression of proteins related to RNA processing and synapse organization (Fig. S1A, D and E), many of which exhibited the same changing trend in *Z^f/f^;N-Cre* brains at P14 (Fig. 1E, Table S1 and S2). Further GO analyses of these overlapping proteins from both datasets again underscored the upregulation of RNA-binding proteins (RBPs) and downregulation of components required for brain development (Fig. 1F, G) in the absence of ZSWIM8. These results not only provide proteome-level evidence supporting our previous findings on the critical role of ZSWIM8 in neurodevelopment, but also suggest a broader requirement for ZSWIM8 in controlling the levels of RBPs.

The fact that RBPs are rich in IDRs, together with our earlier studies on SAX-3, ROBO3 and DAB1 (*24, 29*), led us to hypothesize that ZSWIM8 preferentially recognizes and degrades IDR-containing proteins. Therefore, we utilized three predictor algorithms (VLXT, VSL2b and PrDOS) from the D^2^P^2^ database (*43*) to compare the IDR contents among 3 groups of proteins: 1) those specifically upregulated in ZSWIM8-knockout forebrain at P14, 2) those detected in wild-type forebrain at P14, and 3) the entire annotated mouse proteome (*44*). A common and important theme emerged that, as compared with the reference mouse proteome (*mmu*), IDR-containing proteins (IDR% > 5%) made up a greater proportion of the wild-type forebrain proteome (P14_total) and were even more overrepresented in the upregulated proteins from *Z^f/f^;N-Cre* brains (P14_up) (Fig. 1H). Consistently, the odds of proteins possessing large IDRs (defined as disordered sequences ≥ 30 amino acids long) (*45, 46*) also increased from “*mmu*” to “P14_total” to “P14_up” (Fig. 1I, J). A similar albeit less pronounced trend was seen when the P0 results were used for analysis (Fig. 1K, Fig. S1F and G). These results implicate that IDR-containing proteins in general may have important functions in neo-/post-natal development of the brain, and their particular accumulation in the mutant tissues supports the role of ZSWIM8 as a key regulator of IDR-containing proteins at least in the brain.

### ZSWIM8 regulates miRNA expression in the developing brain

Upregulation of well-known miRNA-binding proteins (miRBPs) such as AGO2 and ELAV1 in absence of ZSWIM8 (Figs. 1A, D and S2B) prompted us to compare our proteomic results with experimentally identified miRNA-binding proteins (miRBPs) from a previous study (*47*). Notably, we found that over a third (65/180) of these miRBPs were upregulated in the *Z^f/f^;N-Cre* brain at P14 (Fig. S2A). Furthermore, small RNA sequencing identified 10 miRNAs consistently upregulated in ZSWIM8 knockout brains (Fig. 2A, B; Table S3).

**Figure 2.**
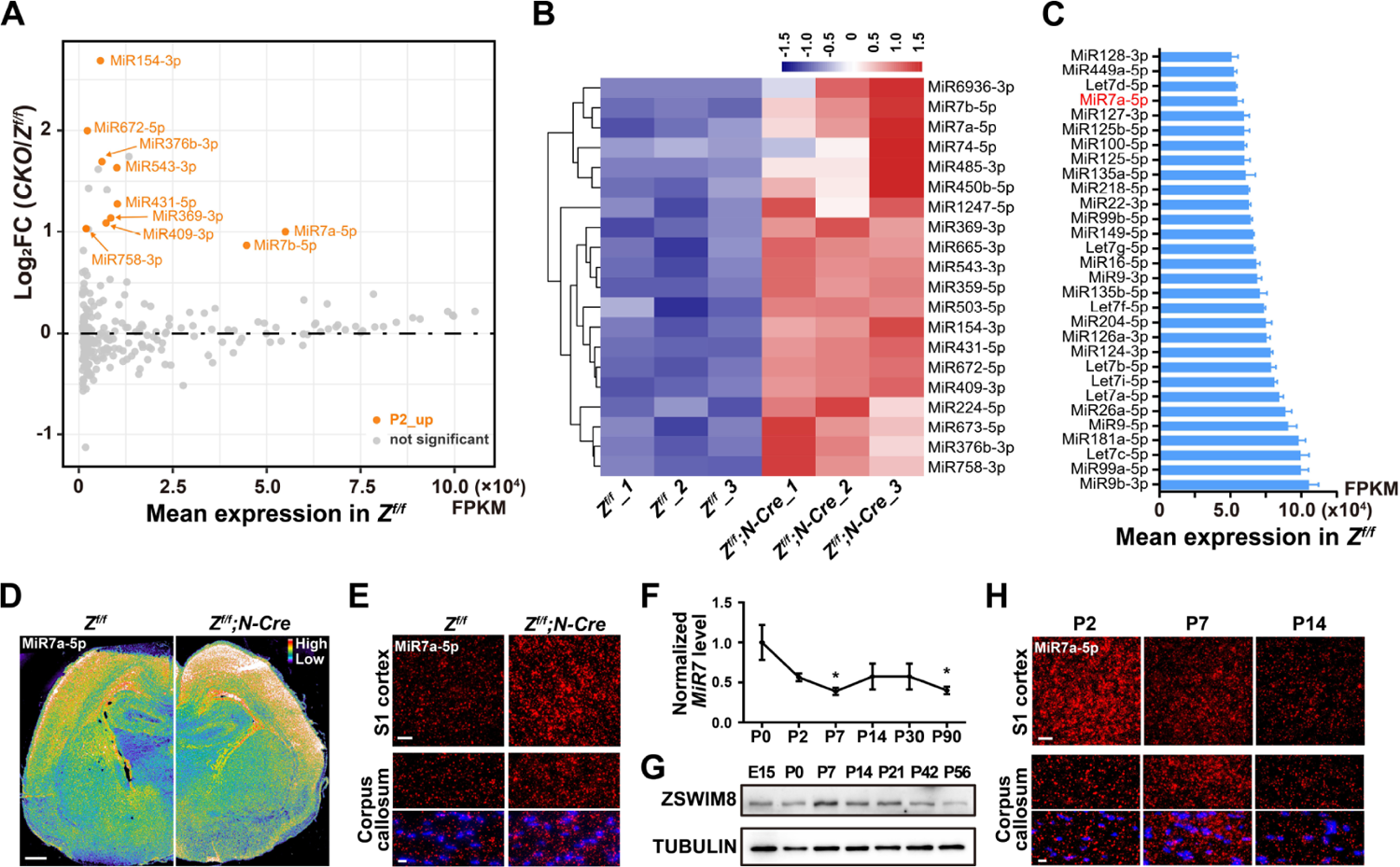
Deletion of ZSWIM8 causes upregulation of a subset of microRNAs in the neonatal mouse brain. **(A)** Whole brain tissues from three pairs of littermates (*Z^f/f^* vs. *Z^f/f^;N-Cre*) at P2 were collected for small RNA-seq analysis. The x-axis represents mean miRNA expression levels in *Z^f/f^* brains. The y-axis represents log_2_-transformed fold-change (FC) of the corresponding miRNAs in *Z^f/f^;N-Cre* brains (CKO). Highlighted in orange are miRNAs with significantly increased expression in *Z^f/f^;N-Cre* animals (p-adjust < 0.05) and a mean expression level > 1000 FPKM in *Z^f/f^*. **(B)** Hierarchic clustering of all miRNAs significantly upregulated in *Z^f/f^;N-Cre* animals (p-adjust <0.05). Data from three pairs of littermates are demonstrated. Blue and orange colors represent low and high expression levels, respectively. (**C**) Mean expression levels of the top 30 most abundant miRNAs in P2 *Z^f/f^* whole brains (N = 3). **(D)** *In situ* visualization of MiR7a in coronal brain sections by miRNAscope from P1 littermates (*Z^f/f^* and *Z^f/f^;N-Cre*) by MiRNAscope. The intensity of signals is illustrated by pseudo color. Scale bar, 100 μm. **(E)** Representative miRNAscope images of MiR7a in the S1 cortex (top, scale bar = 20 μm) and the corpus callosum (bottom, scale bar = 10 μm) of *Z^f/f^* and *Z^f/f^;N-Cre* animals. Nuclei are shown by DAPI staining (blue). **(F)** qRT-PCR quantification of MiR7a levels in whole brains from wild-type (WT) C57BL/6J mice along the indicated time course. Brain tissues from 3 mice were collected for each time point, and MiR7 levels were normalized against the mean at P0. *p < 0.05 relative to P0 (Student’s unpaired *t*-test). **(G)** The protein level of ZSWIM8 in WT forebrain samples at the indicated stages was determined by immunoblotting. E15, embryonic day 15. **(H)** Dynamics of MiR7a expression detected by miRNAscope in the S1 cortex (top, scale bar = 20 μm) and the corpus callosum (bottom, scale bar = 10 μm) of WT mice at the indicated neonatal stages. Nuclei are shown by DAPI staining (blue).

Seven of these (MiR758-3p, MiR376b-3p, MiR409-3p, MiR543-3p, MiR154-3p, MiR369-3p and MiR431-5p) are products of genes clustered at the Chr. 12F1 locus of the mouse genome, implying that their expression may be governed by co-transcription dependent on a ZSWIM8 substrate (Fig. S2C; Table S3). On the other hand, members of the MiR7 family (MiR7a-1, MiR7a-2 and MiR7b) are located on three chromosomes, respectively (Fig. S2C). We examined the spatial and temporal expression of MiR7a, which was among the top 30 most abundant microRNAs in P2 mouse brain (Fig. 2C). Using the miRNAscope technique, we established that MiR7 was widely present in the neonatal brain and highly expressed in the cerebral cortex (Fig. 2D).

Conditional deletion of ZSWIM8 in the embryonic brain by *Nestin*-Driven Cre caused upregulation of overall MiR7 intensity, especially in the S1 cortex and the corpus callosum (Fig. 2D, E). This phenomenon is consistent with the small RNA-Seq results and has also been confirmed by qRT-PCR (Fig. S2D, E). Furthermore, the level of MiR7a was highest at P0, rapidly dropped to less than 50% at P7, and remained largely steady afterwards (Fig. 2F). It is noteworthy that ZSWIM8 protein level in the brain showed an opposite trend from P0 to P7 (Fig. 2G). The miRNAscope data further revealed brain region-specific differences of MiR7a dynamics in the S1 cortex and the corpus callosum at P2, P7 and P14 (Fig. 2H). Such spatiotemporal dynamics supports a regulatory function of MiR7 in brain development, which could be modulated by ZSWIM8.

### ZSWIM8 targets ELAV1 and AGO2 for degradation by different means

ZSWIM8 was first identified in our earlier study as a substrate recognition subunit of the CRL E3 ubiquitin ligase (Fig. 3A, B) (*24*). In this study, we demonstrated that deletion of ZSWIM8 caused dysregulation of numerous proteins involved in RNA metabolism. Thus, we hypothesized that ZSWIM8 probably regulates miRNA expression via controlling the turnover of certain miRBPs. The aforementioned ELAV1 and AGO2 proteins were potential candidates of ZSWIM8 substrates. ELAV1, also known as human antigen R (HuR), has been shown to regulate pri-miRNA processing including that of MiR7 (Fig. S2B) (*48*). Like previously identified substrates such as SAX-3, ROBO3 and DAB1, ELAV1 is a typical IDR-enriched protein based on PONDR prediction (http://www.pondr.com) (Fig. S3A). Indeed, overexpression of wild-type ZSWIM8 evidently enhanced ELAV1 ubiquitination and promoted its degradation in a cycloheximide (CHX) chase assay in HEK293FT cells (Fig. S3B-E). By contrast, the ZSWIM8-ΔBox mutant, which lacks both the BC-box and the CUL2-box and cannot bind the other CRL components (Fig. 3A, B), considerably stabilized ELAV1 (Fig. S3D, E). Under normal conditions, ELAV1 shuttles between the nucleus and cytoplasm (*49*) but is predominantly a nuclear protein (Fig. S3F, G). However, acute sodium arsenite treatment, which induces oxidative stress and protein misfolding (*50, 51*), caused ELAV1 retention in cytosolic stress granules where it nicely colocalized with wild-type ZSWIM8 (Fig. S3F). Interestingly, such colocalization was greatly attenuated in cells expressing the ZSWIM8-ΔIDR mutant (Fig. 3B and Fig. S3G), indicating that the C-terminal IDRs of ZSWIM8 are required for its binding to misfolded ELAV1 under stress conditions. These findings are consistent with the notion that ZSWIM8 uses the “disorder-targets-misorder” mechanism for substrate selection.

**Figure 3.**
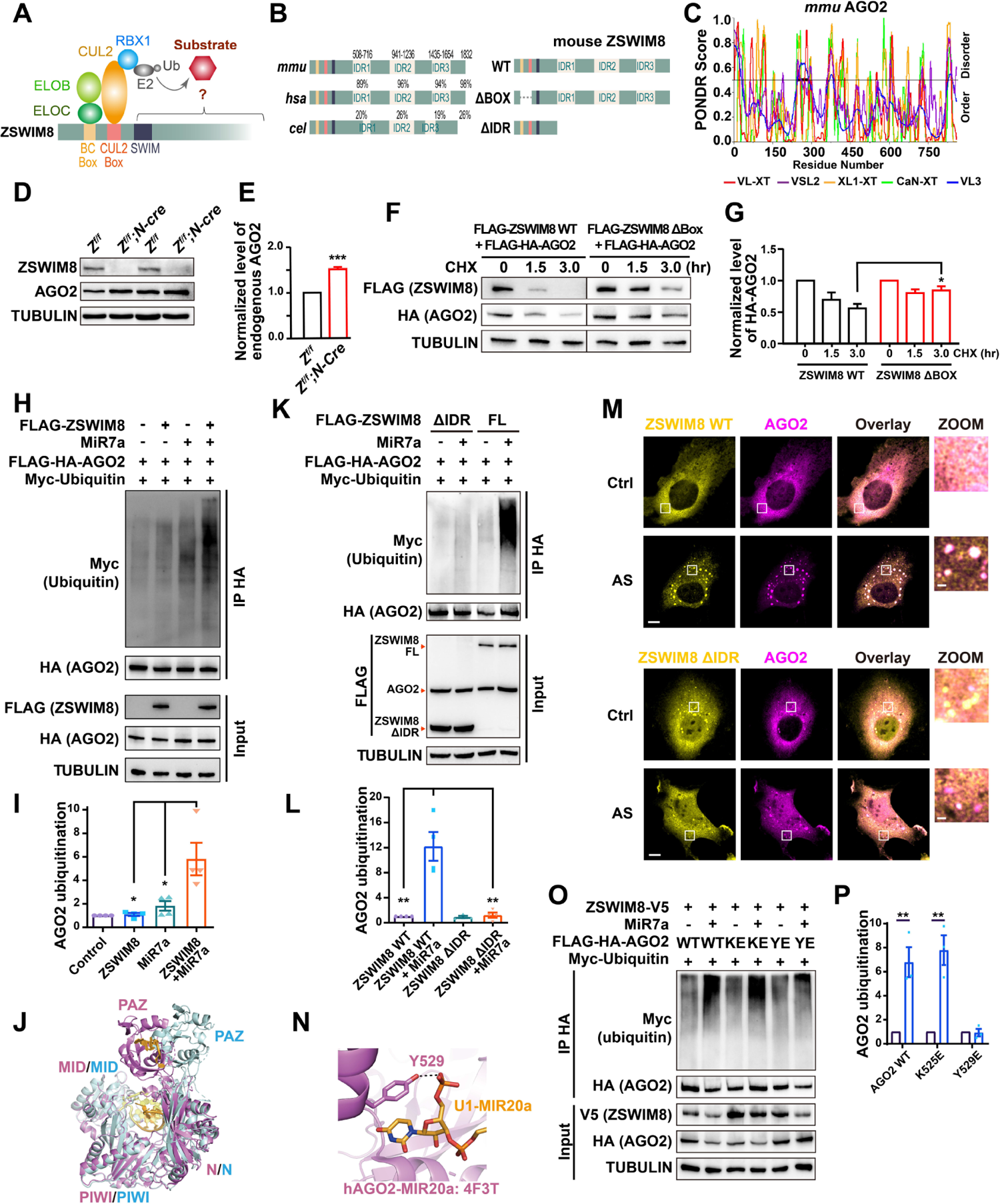
ZSWIM8 promotes AGO2 degradation in an IDR- and miRNA-dependent manner. **(A)** Schematic representation of the CRL^ZSWIM8^ E3 ubiquitin ligase complex. Conserved BC-box, CUL2-box and SWIM motif in the N-terminus of ZSWIM8 are labeled, whereas function of the remainder of ZSWIM8 has not been previously characterized. The BC box and CUL2 box mediate interaction with Elongin B/C and CUL2, respectively. **(B)** Schematic diagrams highlighting the IDRs of ZSWIM8 including their positions and sequence similarities among human, mouse and worm orthologs (left). Truncation mutants of mouse ZSWIM8 used in this study are shown on the right. **(C)** IDR prediction of mouse AGO2 by five algorisms (http://www.pondr.com). The thick black line indicates an IDR only detected by the VL3-BA algorithm, which was trained on both ordered and disordered proteins. This IDR covers 43 amino acid residues (a.a. 247-289) in the PAZ domain of AGO2. **(D)** Immunoblotting of endogenous ZSWIM8 and AGO2 in the forebrain of *Z^f/f^* and *Z^f/f^;N-Cre* animals at P14. Equal amounts of lysates were used for immunoblotting. Tubulin was used as the loading control. **(E)** AGO2 levels as determined in (D) were quantified from 4 pairs of littermates. ***p < 0.001 (Student’s unpaired *t*-test). **(F)** HEK293FT cells were transfected with the indicated constructs and treated with 50 μg / ml cycloheximide (CHX) for 0, 1.5 and 3 h before harvest. Equal amounts of lysates were blotted with anti-FLAG, anti-HA and anti-tubulin antibodies. **(G)** Four independent experiments as in (F) were used for quantification. *p < 0.05 relative to the indicated control (Student’s unpaired *t*-test). **(H)** HEK293FT cells were transfected with the indicated constructs and treated with bortezomib (5 μM, 4 h). FLAG-HA-AGO2 was immunoprecipitated by anti-HA nanobody beads, and polyubiquitination signals were detected by anti-MYC antibodies. **(I)** Four independent experiments as in (H) were used for quantification. *p < 0.05 relative to the indicated control (one-way ANOVA). **(J)** Structural superposition of the hAGO2-MIR20a complex (PDB: 4F3T, pink) and hAGO2-MIR27a bound to target RNA (PDB: 6MFN, blue) showing considerable flexibility of the PAZ domain upon mRNA binding. **(K)** Polyubiquitination of FLAG-HA-AGO2 in the presence of FLAG-ZSWIM8 WT or FLAG-ZSWIM8 ΔIDR were examined as in (H). **(L)** Four independent experiments as in (J) were used for quantification. *p < 0.01 and **p < 0.01 relative to the indicated control (one-way ANOVA). **(M)** Immunostaining images of V5-tagged ZSWIM8 (WT or ΔIDR) and HA-tagged AGO2 in HUVEC cells. Scale bar, 10 μm. Sodium arsenite (AS, 500 μM, 30 min) was used to induce oxidative stress and formation of cytosolic granules. Zoom-in images of the boxed regions are shown on the right. Scale bar, 2 μm. **(N)** Structural basis for the critical role of AGO2-Y529 in miRNA binding (PDB: 4F3T). The highly conserved Y529 residue locates in the miRNA-binding pocket of AGO2. The Y529E mutation that mimics phosphorylation at this site strongly inhibits loading of miRNA into AGO2. **(O)** HEK293FT cells were transfected as indicated constructs and treated with bortezomib (5 μM, 4 h). FLAG-HA-AGO2 was immunoprecipitated by anti-HA nanobody beads. Polyubiquitination of AGO2 variants was determined by anti-MYC antibodies. KE: K525E; YE: Y529E. **(P)** AGO2 ubiquitination as in (O) was quantified from 3 independent experiments. **p < 0.01 (Student’s unpaired *t*-test).

AGO2 has been suggested to be a substrate of ZSWIM8 involved in TDMD of MiR7 in various cell lines (*33, 34*). We also found that AGO2 was consistently upregulated in *Z^f/f^;N-Cre* brain tissues (Fig. 3D, E). However, different from IDR-enriched ELAV1, AGO2 is organized into several well folded domains (*52*) and only the PAZ domain shows weakly disorder characteristics (Fig. 3C). Direct evidence of ZSWIM8-mediated AGO2 degradation and the molecular details thereof have been lacking. Here, we found that the BC-box and CUL-2 box of ZSWIM8 were required for AGO2 degradation in HEK293FT cells (Fig. 3F, G), as seen with ELAV1 (Fig. S3D, E). However, simple overexpression of wild-type ZSWIM8 only marginally increased AGO2 ubiquitination (Fig. 3H, I), which was distinct from the case of ELAV1 (Fig. S3B, C). Structural comparison of AGO2 in its miRNA-bound versus miRNA/mRNA-bound forms showed considerable movement of the PAZ domain (*52–54*) (Fig. 3J). Thus, we hypothesized that RNA binding may trigger conformational distortion of AGO2 particularly in the PAZ domain, and provide structural features that can be recognized by ZSWIM8 for ubiquitination.

To test this possibility, we first overexpressed MiR7a together with wild-type ZSWIM8 and surprisingly found that excessive MiR7a strongly stimulated ZSWIM8-mediated AGO2 ubiquitination (Fig. 3H, I). Moreover, this effect was dependent on the IDRs of ZSWIM8 (Fig. 3K, L), the lack of which not only inhibited the activity of ZSWIM8 towards AGO2 but also greatly attenuated colocalization of ZSWIM8 and AGO2 within arsenite-induced cytoplasmic granules (Fig. 3M). Then we reasoned that RNA binding may be crucial for AGO2 regulation, given its role in RNA interference and TDMD. To examine this hypothesis, we took advantage of two RNA-binding mutants of AGO2, K525E and Y529E (Fig. 3N). The AGO2 Y529E mutant, which cannot bind miRNA (*55, 56*), failed to be ubiquitinated by ZSWIM8 beyond the background level, even in the presence of MiR7 (Fig. 3 O, P). The K525E mutation, which severely impairs target mRNA binding but without affecting miRNA association (*57*), had no discernable effects on AGO2 ubiquitination by ZSWIM8 (Fig. 3O, P). Therefore, we conclude that association of miRNA (e.g., MiR7) is indeed a prerequisite for AGO2 ubiquitination and degradation, which in turn deprotects MiR7. These results represent an unprecedented, miRNA-dependent mechanism of substrate recognition by quality-control E3s such as ZSWIM8.

### ZSWIM8 deletion in nervous system impairs oligodendrocyte maturation and myelination

We noticed from the proteomic results that deletion of ZSWIM8 caused significant downregulation of myelination-related proteins, such as CNP, MBP and MYPR (Fig. 1A), implicating previously unknown functions of ZSWIM8 in oligodendrocyte lineage cells. Oligodendrocytes (OLs) are specialized glial cells that form myelin sheath around nerve fibers in the brain and provide electrical insulation to increase axonal conduction velocity (*58*).

Oligodendrocyte progenitor cells (OPCs) start to appear at the late embryonic stage and differentiate into oligodendrocytes (OLs) after birth (*59–61*). In the postnatal forebrains, the level of OPC-specific marker PDGFRA reached a peak at P7 (*62*). Afterwards, the expression of PDGFRA quickly dropped, accompanied by gradual increase of markers for premyelinating and mature OL markers (CNP and MBP, respectively) (Fig. 4A). At P14, levels of OPC-specific PDGFRA and the general oligodendrocyte-lineage marker OLIG2 showed no significant difference between the *Z^f/f^* and *Z^f/f^;N-Cre* forebrains (Fig. 4B, C). However, the levels of both CNP and MBP were markedly reduced upon deletion of ZSWIM8 (Fig. 4B, C). Consistently, anti-MBP immunohistochemistry (IHC) staining showed severe thinning of the corpus callosum, a connective tissue largely formed by myelinated axons and oligodendrocyte lineage cells, in P14 *Z^f/f^;N-Cre* brains (Fig. 4D, F). Myelinated areas were also significantly reduced within this structure (Fig. 4E, G). A similar defect in myelin sheath was seen in the S1 cortex of *Zswim8-*null brains (Fig. S4A-C). These data together suggest that absence of ZSWIM8 from the neonatal brain may causes impaired oligodendrocyte maturation and function.

**Figure 4.**
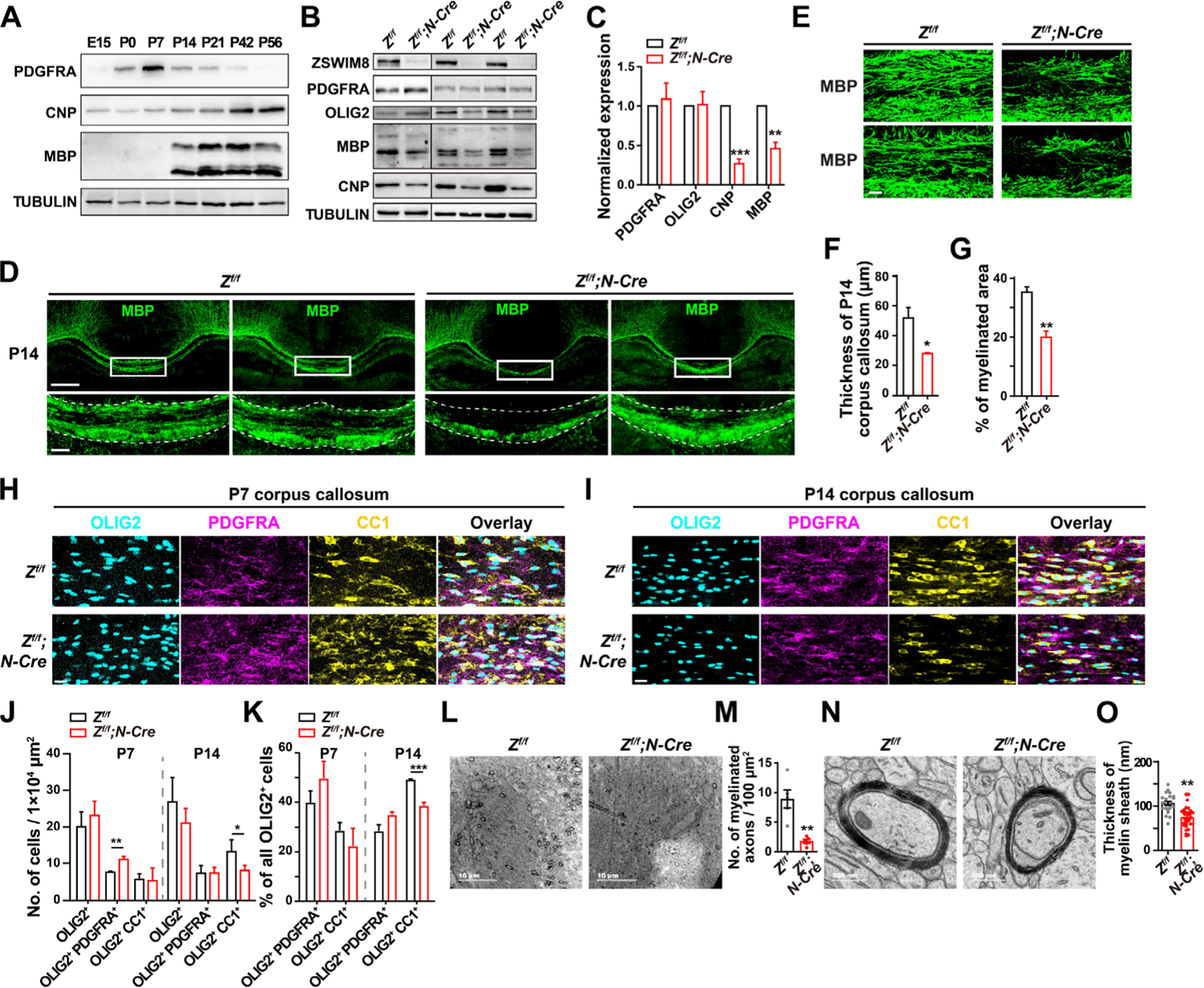
ZSWIM8 is required for oligodendrocyte maturation and myelination. **(A)** Western blot analysis of PDGFRA (OPC marker), CNP (premyelinating OL marker) and MBP (mature OL marker) in the forebrain from WT mice of the indicated ages. **(B)** Forebrain samples were collected from 3 pairs of littermates (*Z^f/f^* and *Z^f/f^;N-Cre*) at P14 and probed with the indicated antibodies. OLIG2 is a marker for oligodendrocyte lineage cells. **(C)** Quantification of protein levels shown in (B). **p < 0.01 and ***p < 0.001 (Student’s unpaired *t*-test). **(D)** Immunohistochemistry (IHC) of MBP in coronal brain sections from *Z^f/f^* and *Z^f/f^;N-Cre* littermates (P14). Scale bar, 100 μm (top). Magnified images of the corpus callosum in the boxed medial regions are shown at the bottom (scale bar = 20 μm). The corpus callosum is outlined by dotted lines based on DAPI staining (not shown). **(E)** High magnification images (60 X) of anti-MBP IHC showing myelinated areas in the corpus callosum. Scale bar, 20 μm. **(F)** Quantification of the thickness of the corpus callosum (marked by white dotted lines in D). N = 3 pairs of littermates. *p < 0.05 (Student’s unpaired *t*-test). **(G)** Quantification of the myelinated area in the corpus callosum based on MBP intensities in (E). N = 3 pairs of littermates. **p < 0.01 (Student’s unpaired *t*-test). **(H, I)** Corpus callosum tissues from *Z^f/f^* and *Z^f/f^;N-Cre* animals at P7 (H) and P14 (I) were immunostained for OLIG2 (OL lineage marker), PDGFRA (OPC marker) and CC1 (mature OL marker). Scale bar, 20 μm. **(J)** The density of OLIG2-marked OL lineage cells, OLIG2^+^ PDGFRA^+^ OPCs and OLIG2^+^ CC1^+^ mature OLs in the corpus callosum was quantified in *Z^f/f^* and *Z^f/f^;N-Cre* animals at indicated stages. N = 3 pairs of littermates. *p < 0.05 and **p < 0.01 (Student’s unpaired *t*-test). **(K)** The percentage of OPCs or mature OLs among all OLIG2^+^ cells in the corpus callosum were quantified for each condition. N = 3 pairs of littermates. ***p < 0.001 (Student’s unpaired *t*-test). **(L)** Representative transmission electron microscopy (TEM) micrographs of the corpus callosum from P14 *Z^f/f^* and *Z^f/f^;N-Cre* animals are demonstrated. Scale bar, 10 μm. **(M)** The numbers of myelinated axons in random 100 μm^2^ areas (N = 5) were quantified in (L) for each littermate. **p < 0.01 (Student’s unpaired *t*-test). **(N)** Representative transmission electron microscopy (TEM) images of cross sections of single axons from P14 *Z^f/f^* and *Z^f/f^;N-Cre* animals are demonstrated. Scale bar, 500 nm. **(O)** Quantification of the thickness of myelin sheath on individual axons (N = 20∼30). **p < 0.01 (Student’s unpaired *t*-test).

Next, we examined the effects of ZSWIM8 deletion on oligodendrocyte development. At P7, OPCs marked by PDGFRA/OLIG2 double-positive staining (OLIG2^+^ PDGFRA^+^) actually accumulated at a higher density in the corpus callosum of *Z^f/f^;N-Cre* mice than in that of *Z^f/f^* controls (Fig. 4H, J and K). The numbers of OLIG2^+^ CC1^+^ mature OLs were low in both groups at this stage. At P14, an increase of mature OLs was observed in *Z^f/f^* controls, but deletion of ZSWIM8 abolished this change (Fig. 4I-K). Total OLIG2^+^ cells remained comparable in the number and density between controls and KO samples at both P7 and P14 (Fig. 4H-K). A similar reduction in mature OLs was observed in deep layers of the S1 cortex in *Z^f/f^;N-Cre* mice (Fig. S4D-F).

We further examined ultrastructural changes in myelination in P14 corpus callosum using transmission electron microscopy (TEM). Both the number of myelinated axons (Fig. 4L-M) and the thickness of myelin sheath (Fig. 4N-O) were significantly reduced in *Z^f/f^;N-Cre* mutant brains. Together, our results from multiple assays indicate that deletion of ZSWIM8 impedes oligodendrocyte maturation, leading to hypomyelination in neonatal brains.

### ZSWIM8 ablation in the oligodendrocyte lineage leads to hypomyelination

To further determine whether ZSWIM8 regulates myelination through cell-autonomous mechanisms, we examined expression and requirement of ZSWIM8 in oligodendrocyte lineage cells. In the corpus callosum and S1 cortex of P14 *Zswim8^f/f^* mice, *Zswim8* mRNA was detected in both *Pdgfra^+^* and *Cnp^+^* cells by RNAscope (Fig. 5A and Fig. S5A), but was significantly more abundant in the latter (Fig. 5B). Next, we generated a *Zswim8^f/f^; Cnp-Cre* strain (*Z^f/f^; C-Cre*) in which ZSWIM8 was specifically eliminated in premyelinating oligodendrocytes (pre-OLs) when the *Cnp* promoter started to express. Reduced ZSWIM8 expression in *CNP^+^* but not in *Pdgfra^+^* cells confirmed specific knockout of ZSWIM8 in differentiating oligodendrocytes (Fig. 5A and Fig. S5A). Notably, *Cnp* mRNA drastically decreased in *Z^f/f^;C-Cre* mutants compared to *Z^f/f^*, consistent with downregulation of CNP proteins in the forebrain of P14 *Z^f/f^;N-Cre* mutant animals (Fig. 5C). Additionally, in agreement with results from *Z^f/f^;N-Cre* mice, anti-MBP IHC showed clear hypomyelination in the corpus callosum and S1 cortex of *Z^f/f^;C-Cre* animals at P14 (Fig. 5D and Fig. S5B). Both the myelinated area in the corpus callosum (Fig. 5E) and the length of myelin sheath in S1 deep layers (Fig. S5C-D) significantly decreased in *Z^f/f^;C-Cre* brains. These data corroborate that ZSWIM8 is essential for maturation of oligodendrocytes and formation of myelin sheath in a cell-autonomous fashion.

**Figure 5.**
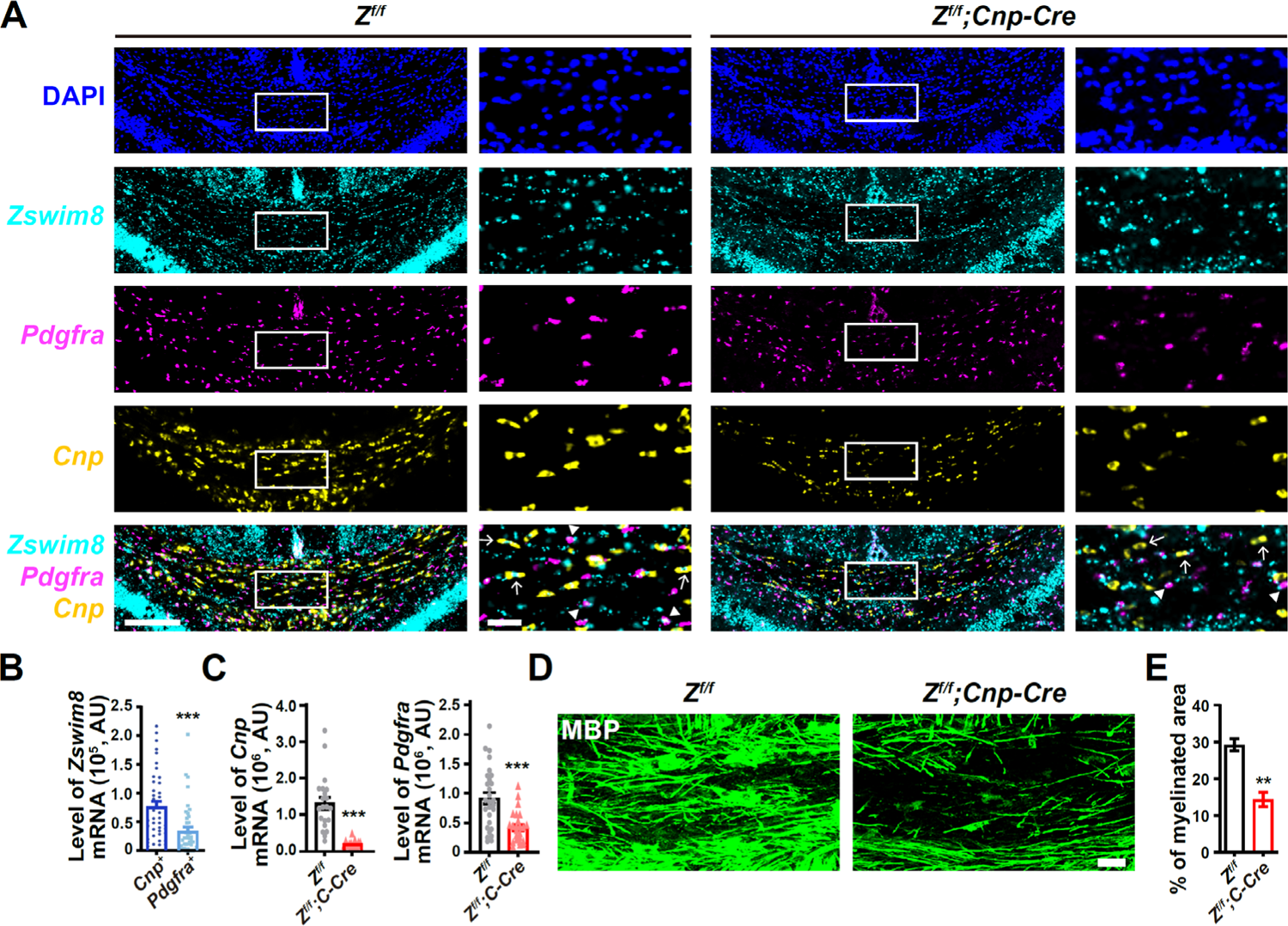
Conditional deletion of ZSWIM8 in premyelinating OLs impairs OL maturation and causes hypomyelination in the corpus callosum. **(A)** Representative RNAscope images of *Zswim8*, *Pdgfra* and *Cnp* in the corpus callosum (coronal sections) of P14 *Z^f/f^* and *Z^f/f^;C-Cre* mice. Scale bar, 50 μm. Zoom-in images of the boxed regions are shown on the right (scale bar = 10 μm). White arrowheads and arrows point to *Pdgfra*^+^ or *Cnp^+^* cells that co-expressed *Zswim8*, respectively. **(B)** Integrated cellular levels of *Zswim8* mRNA (AU, arbitrary unit) detected by RNAscope in *Cnp^+^* pre-OLs and *Pdgfra^+^* OPCs within the corpus callosum of P14 *Z^f/f^* mice (N = 40 cells). ***p < 0.001 (Student’s unpaired *t*-test). **(C)** Integrated cellular levels of *Cnp* and *Pdgfra* mRNA detected by RNAscope in the corpus callosum from P14 *Z^f/f^* and *Z^f/f^;C-Cre* mice in (A) were quantified (N = 40 cells). ***p < 0.001 (Student’s unpaired *t*-test). **(D, E)** Immunohistochemistry images of MBP at the midline of the corpus callosum (coronal sections) from P14 *Z^f/f^* and *Z^f/f^;C-Cre* animals. Scale bar, 20 μm. Quantification of the myelinated areas is shown in (E). N = 3 pairs of littermates. **p < 0.01 (Student’s unpaired *t*-test).

### MiR7 agomir inhibits oligodendrocyte maturation and function

A previous *in vitro* study suggested involvement of MiR7 in oligodendrocyte differentiation, the level of which must be tightly controlled (*42*). We wondered whether impaired OL development and hypomyelination in ZSWIM8-deleted mice could be ascribed to MiR7 dysregulation. MicroRNAscope and immunohistochemistry assays confirmed that MiR7 was expressed in OLIG2^+^ oligodendrocytes lineage cells in the corpus callosum at neonatal stages (Fig. 6A). To address whether MiR-7 acts downstream of ZSWIM8 *in vivo*, we first used MiR7 agomir, a chemically modified double-stranded RNA that can mimic the function of endogenous MiR7. Compared to common miRNA mimics, MiR7 agomir shows higher affinity for cell membrane and better stability and is particularly suitable for *in vivo* experiments. We first injected MiR7 agomir into bilateral ventricles in wild-type brains at P3, when endogenous MiR7 drastically drops, and then examined OPCs and OLs by IHC after a week (Fig. 6B). In the corpus callosum, total OLIG2^+^ cells and OLIG2^+^ PDGFRA^+^ OPCs showed no difference between control and MiR7 agomir-treated animals (Fig. 6C-E). Consistent with observations in ZSWIM8-deleted animals, the density and percentage of OLIG2^+^ CC1^+^ mature OLs were significantly reduced by MiR7 agomir, pointing to an inhibitory function of MiR7 in oligodendrocyte maturation (Fig. 6C-E). In addition, downregulation of CNP and MBP proteins in the corpus callosum was also observed by in MiR7 agomir-treated animals (Fig. S6A). Intriguingly, both human and mouse *Cnp* 3’UTRs contain a predicted MiR7 targeting site (Fig S6B). Together, these data indicate that decline of the MiR7 level is necessary for oligodendrocyte maturation in the neonatal mouse brain by allowing the expression of key myelination factors such as CNP.

**Figure 6.**
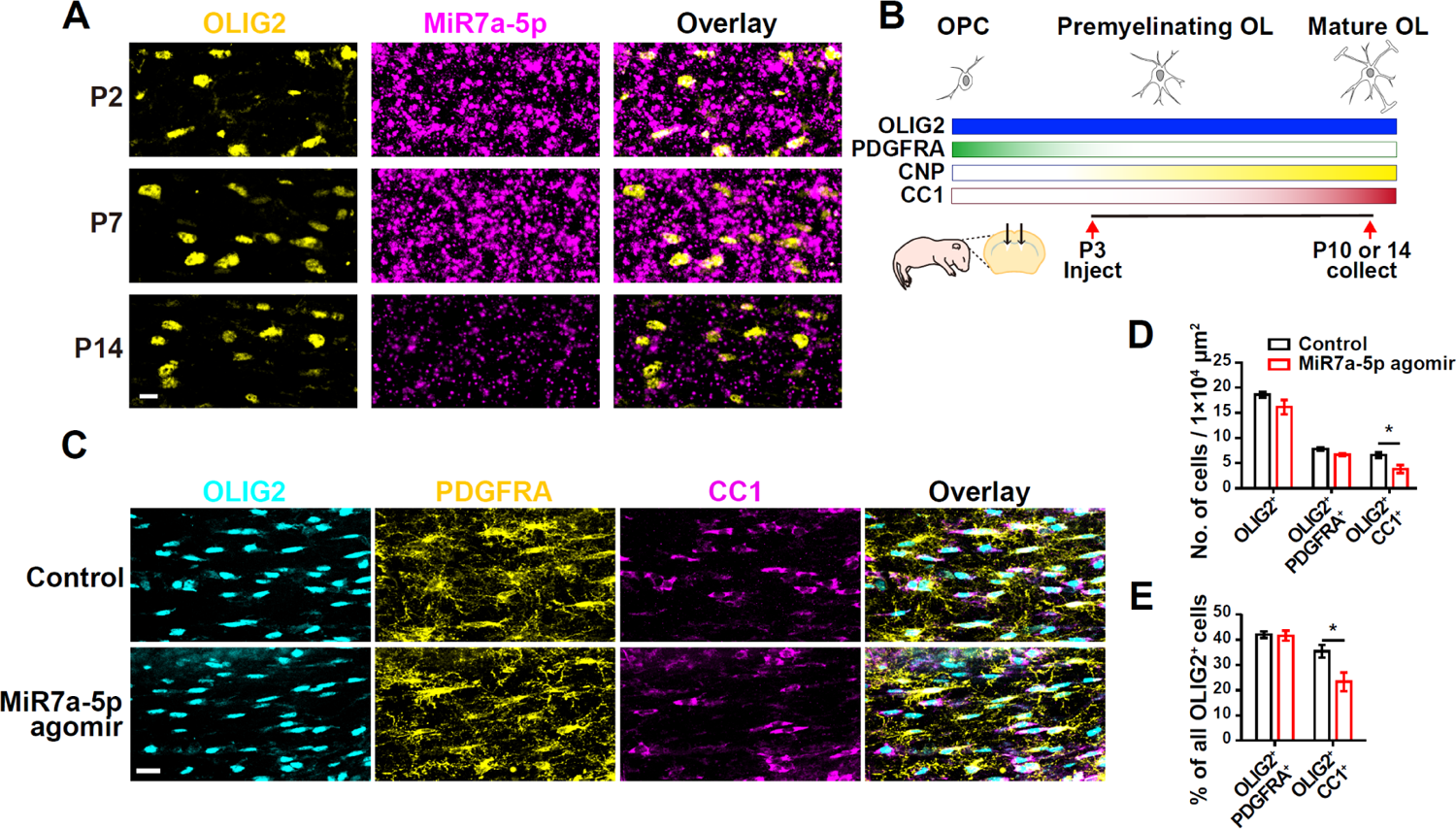
Excessive MiR7 inhibits oligodendrocyte maturation and CNP expression *in vivo*. **(A)** miRNAscope detection of MiR7 in OLIG2^+^ cells in the corpus callosum from WT mice at neonatal stages. Scale bar, 20 μm. **(B)** Illustration of the expression profiles of oligodendrocyte lineage markers during development. The experimental design of *in vivo* injections of MiR7 agomir is shown at the bottom. **(C)** WT brains that received control or MiR7a agomir were analyzed on P10 for OLIG2, PDGFRA and CC1 expression in the corpus callosum by immunohistochemistry. Scale bar, 20 μm. **(D)** The density of OLIG2^+^ cells, OLIG2^+^ PDGFRA^+^ OPCs and OLIG2^+^ CC1^+^ mature OLs in the corpus callosum from mice treated with control or MiR7a agomir were quantified. *p < 0.05 (Student’s unpaired *t*-test). **(E)** The percentage of OPCs or mature OLs among all OLIG2^+^ cells in the corpus callosum from mice treated with control or MiR7a agomir were quantified. *p < 0.05 (Student’s unpaired *t*-test).

### ZSWIM8 regulates oligodendrocyte development through MiR7

Finally, we performed loss-of-function experiments to establish epistasis between ZSWIM8 and MiR7 in regulation of myelination. To this end, we injected MiR7 antagomir, a chemically modified single-strand miRNA inhibitor which suppresses endogenous MiR7 activity, into bilateral ventricles of *Z^f/f^* and *Z^f/f^;C-Cre* mice at P3 following the scheme shown in Fig. 6B. Maturation of OLs were then examined at P14 by OLIG2 and CC1 staining. In the control-treated groups, *Z^f/f^;C-Cre* mutant brains showed a decrease in the thickness of corpus callosum and in the percentage of OLIG2^+^ CC1^+^ mature OLs as compared to *Z^f/f^* (Fig. 7A, B and D), recapitulating our earlier observations (Fig. 4), although the density of mature OLs in these animals only showed a subtle decrease (Fig. 7C). In *Z^f/f^;C-Cre* mice receiving injections of MiR7 antagomir, however, the thickness of the corpus callosum was restored to the level of control animals (Fig. 7B). The density and percentage of mature OLs were also significantly increased compared to control-treated *Z^f/f^;C-Cre* animals (Fig. 7A-D). RNAscope assays also showed that the weakened expression of *Cnp* in *Z^f/f^; C-Cre* mice was elevated by MiR7 antagomir (Fig. S7A). Meanwhile, we noticed that MiR7 antagomir mildly reduced the corpus callosum thickness and increased the total OLIG2^+^ cells in *Z^f/f^* controls (Fig. 7B, C), suggesting that the activity of MiR7 must be fine-tuned to an optimal level during the neonatal stage. Altogether, these data demonstrate that MiR7 functions downstream of ZSWIM8 to control oligodendrocyte development *in vivo*. The coordinated action of ZSWIM8, AGO2 and MiR7 in OL maturation and functions (Fig. 7E) has illustrated the biological importance of IDR-directed protein quality control and microRNA metabolism in the developing CNS. A better understanding of the operating principles of such systems will bring new insights into the pathogenesis and treatment of related diseases.

**Figure 7.**
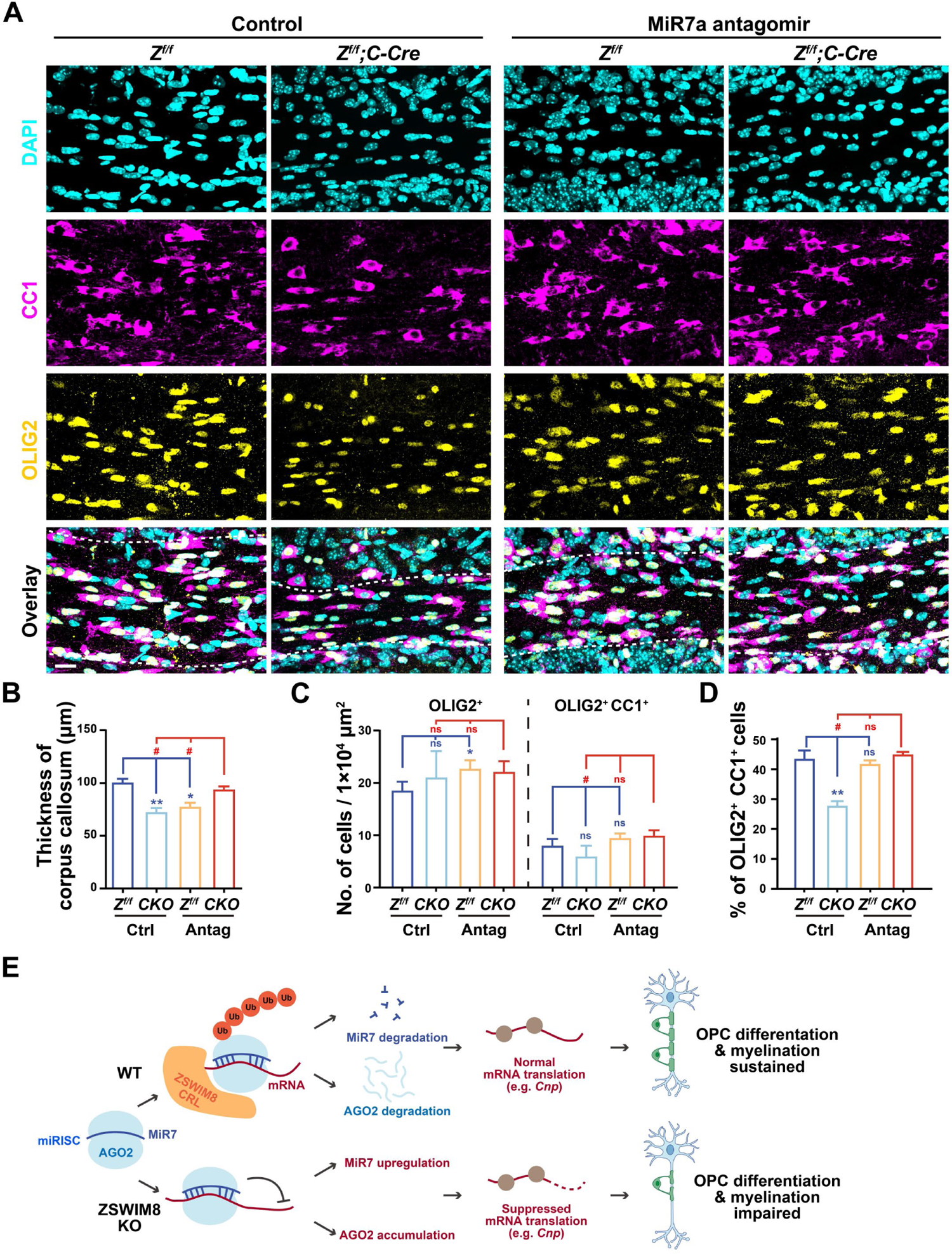
MiR7 antagomir rescues oligodendrocyte maturation in ZSWIM8 knockout brains. **(A)** Immunohistochemistry of OLIG2 and CC1 in the corpus callosum (coronal sections) from P14 *Z^f/f^* and *Z^f/f^;C-Cre* brains that received control or MiR7 antagomir on postnatal day 3 (P3). DAPI stains the nuclei. The corpus callosum is demarcated by white dotted lines. Scale bar, 20 μm. **(B)** Quantification of the thickness of corpus callosum in (A). N = 3 for each group. *p < 0.05, **p < 0.01, and ^#^p < 0.05 (one-way ANOVA) relative to indicated samples. **(C)** Densities of OLIG2^+^ and OLIG2^+^ CC1^+^ cells in the P14 corpus callosum of the indicated groups were quantified. *p < 0.05 and ^#^p < 0.05 (one-way ANOVA) relative to indicated samples. ns, not significant. **(D)** The percentage of OLIG2^+^ CC1^+^ mature OLs in all OLIG2^+^ cells was quantified (one-way ANOVA). **p < 0.01 and ^#^p < 0.05 (one-way ANOVA) relative to indicated samples. ns, not significant. **(E)** A schematic model (created with BioRender, https://www.biorender.com) demonstrating how ZSWIM8 controls oligodendrocyte differentiation and myelination through its regulation of AGO2 degradation and microRNA metabolism.

## Discussion

The pathophysiological importance of IDRs of proteins has become widely recognized, and our observation from the current study supports the biological relevance of IDR-proteins in cells undergoing drastic proteomic reshaping, such as during brain development. These proteins, however, pose a formidable challenge to the quality control system in cells since their disordered nature defies clearance by effectors that recognize finite structures. Rather, a fuzzy mode of substrate recognition has been proposed for quality control enzymes such as the San1 E3 ubiquitin ligase and the Hsp90 molecular chaperone to cope with a wide range of targets/clients with unstructured sequences (*20, 25*). Our work in mice and human cells demonstrate that this “disorder-targets-misorder” mechanism is conserved in mammals, and that CRL^ZSWIM8^ is one of the first mammalian E3s known to utilize this mechanism for broad regulation of IDR-containing proteins.

ZSWIM8 is a large protein of over 1,800 amino acids in length. Except for the N-terminal domains, the remainder of this protein is poorly annotated in terms of structure and function. And yet the number, sizes and spacing of large IDRs in this region are highly conserved from worm to human, strongly suggesting functional importance (Fig. 3B) (*24*). Deletion of these IDRs abolished the activity of ZSWIM8 toward its substrates, in accordance with the “disorder-targets-misorder” notion. It will be interesting to determine the specific contributions of each individual IDR in substrate recognition in the future. On the other hand, we noticed that wild-type ZSWIM8 itself is relatively unstable, and deletion of either IDRs or the BC Box led to appreciable stabilization of the protein (Fig. 3F, J). This suggests that ZSWIM8 could auto-ubiquitinate and regulate its own level, considering that itself is an IDR-rich protein. Alternatively, ZSWIM8 might be targeted by other IDR-directed E3(s), the identity of which would be very interesting to find out.

During the preparation of this manuscript, two new studies (*31, 32*) reported perinatal lethality of global knockout of ZSWIM8 in mice, confirming our earlier results (*29*). These investigators identified developmental defects in the heart and lungs of the mutant animals, and systematically analyzed changes in miRNA profiles caused by ZSWIM8 deletion. From a different route focusing on IDR protein degradation in neonatal brains, we arrived at the same conclusion that ZSWIM8 monitors miRBPs and finely controls the expression and function of their associated miRNAs (e.g., MiR7). All of the upregulated miRNAs we found in the *Zswim8^f/f^;N-Cre* brain were also shown to be increased in whole-body ZSWIM8 knockouts (*31, 32*). These results, together with other recent findings from worm, fly and human cell culture systems (*30, 63–66*) further corroborate the physiological relevance of ZSWIM8 in regulating miRNAs and miRBPs.

Although both ELAV1 and AGO2 regulate MiR7 and can be ubiquitinated by ZSWIM8, they differ in the molecular details. As a typical IDR-protein, ELAV1 recognition and ubiquitination by ZSWIM8 shares a similar mechanism as seen previously with the other ZSWIM8/EBAX-1 substrates (*24, 29*). ELAV1 antagonizes processing of miRNAs including MiR7 (*67, 68*). ZSWIM8 selectively targets misfolded ELAV1 for degradation to protect activity of properly folded ELAV1 toward miRNA. In ZSWIM8-null tissues, normal ELAV1 proteins are either functionally interfered or diluted by misfolded ELAV1, thus insufficient to keep the level of miR7 under control. Distinct from ELAV1, AGO2 is a rather structured protein with weakly disordered properties in the PAZ domain (Fig. 3C). Its ubiquitination by ZSWIM8 requires the participation of miRNA as we demonstrated by MiR7 overexpression and the miRNA binding-deficient AGO2 mutant (Fig. 3). This finding adds a new piece of evidence that RNAs, including miRNA shown here and previously reported circular RNAs (circRNAs) and long non-coding RNAs (lncRNAs) (*30, 69–71*), can act as important mediators of RBP degradation by the UPS. One possibility is that miRNAs might be directly involved in substrate/enzyme recruitment as circRNAs and lncRNAs. It is also possible that MiR7-stimulated AGO2 degradation probably involves conformational change of AGO2 that exposes cryptic binding sites and/or ubiquitination sites favored by the ZSWIM8 ligase (Fig. S7B). We also cannot rule out the possibility that other entities of the AGO-miRNA complex (e.g., other RBPs) in the cell may also contribute to ZSWIM8 binding.

AGO2 and miRNAs form the RISC complex to downregulate target mRNA expression, while ZSWIM8 degrades AGO2 and consequentially deprotects the bound miRNAs, leading to TDMD (*31–34*). Loss of ZSWIM8 disrupts this self-limiting system, thus leads to aberrant expression of miRNAs such as MiR7. Multiple studies have reported involvement of MiR7 in brain development and neurological diseases (*38*). As one of the most abundant miRNAs in the brain, MiR7 functions in various brain regions regulating the expression of several protein-coding and non-coding RNAs (*38, 72*). Our work presented the first *in vivo* evidence that ZSWIM8-dependent downregulation of MiR7 in OPCs is essential for OL maturation and functions at neonatal stages. These results not only highlight the importance of spatiotemporal and cell-type specificity of miRNA function and regulation, but also set the stage for future investigations on quality control of IDR-containing proteins by ZSWIM8 in health and disease.

## Materials and Methods

### Mice

*Zswim8^flox/flox^* mice previously described (*29*) were crossed with *Nestin-Cre* (*73*) or with *Cnp-Cre* mice (*74, 75*) to generate conditional knockouts. All mice were maintained at the Laboratory Animal Center at Zhejiang University, follwing the policies of the Institutional Animal Core and Use Committee (IACUC) and approved animal protocols.

### Plasmids

pcDNA6b-FLAG-mZSWIM8-V5 WT and ΔBox constructs have been reported (*29*). The ΔIDR mutant was made from pcDNA6b-FLAG-mZSWIM8-V5 WT by deleting aa. 508-1830 of mZSWIM8. Pri-MiR7a sequence was synthesized and inserted into the pCMV-MCS-EF1a-copGFP vector between the EcoRI and BamHI sites. ELAV1 was PCR-amplified from the Ultimate^TM^ ORF Clone Collection (Thermo Fisher Scientific) and inserted into the pcDNA3.1-HA-gtwy vector (pCZGY58) by Gibson assembly. FLAG-HA-AGO2 was kindly provided by Dr. Dongli Pan (Zhejiang University). Site-directed mutagenesis was performed to obtain the AGO2-K525E and Y529E. pBOBI-Myc-ubiquitin was a gift from Dr. Ying Liu (Peking University). All plasmids have been confirmed by Sanger sequencing.

### Cell culture, immunocytochemistry and biochemistry

HEK293FT and HUVEC cells were grown in DMEM (Thermo Fisher Scientific) supplemented with 10% fetal bovine serum (Gibco) and 1% Penicillin/Streptomycin (Yeasen). Transfection was carried out with Lipofectamine 8000 (Beyotime).

For immunocytochemistry, HUVEC cells grown on coverslips were transfected with indicated constructs for 24 h. To induce cellular stress and formation of cytosolic granules, cells were treated with 500 μM sodium arsenite for 30 min and then immediately fixed with 4% paraformaldehyde/4% sucrose in PBS for 20 min. Cells were then permeabilized with 0.2% Triton X-100 in PBS for 5 min. After rehydration in PBS, cells were pre-blocked with 3% BSA in PBS, and then incubated with primary antibodies at 4°C overnight followed by fluorescent secondary antibodies for 1 h at room temperature (RT) as previously described (*29*). Images were acquired using an Olympus spinning disc confocal microscope with a 100 X objective and analyzed with MetaMorph (Universal Imaging Corporation).

For pulse-chase assays, HEK293FT cells were transfected with indicated constructs. Next day, cells were treated with 50 μg / ml cycloheximide (CHX) for indicated time. Twenty-four hours after transfection, cells were collected in cold TENT buffer (50 mM Tris, pH 7.4, 1 mM EDTA, 150 mM NaCl, 0.5 % Triton X-100) supplemented with protease inhibitors on ice. After centrifugation at 14,000 rpm for 10 min, supernatants were saved for concentration measurement and then processed for SDS-PAGE and western blotting.

To detect ubiquitination, HEK293FT cells transfected with indicated constructs. Then cells were treated with 5 µM bortezomib for 4 h on next day and then lysed in TENT buffer supplemented with 1% SDS, protease inhibitors and 20 mM N-ethylmaleimide. After further denaturation at 95°C for 5 min, cell lysates were diluted 10-fold with regular TENT buffer without SDS and needle-sheared for 20 times. Then these lysates were cleared by centrifugation at 14,000 rpm for 10 min. The supernatants were immunoprecipitated with 10 μl of anti-HA nanobody agarose beads (AlpaLifeBio) at 4°C for 2 h. Beads were washed 5 times with TENT buffer (supplemented with 0.1% SDS and protease inhibitors) and then processed for SDS-PAGE and western blotting.

Whole brain or forebrain tissues from control and mutant mice at indicated ages were collected and homogenized in TENT buffer supplemented with protease inhibitors. The supernatants after clarification by centrifugation (14,000 rpm, 30 min, 4°C) were collected for SDS-PAGE and immunoblotted for indicated antibodies.

For all biochemistry assays, protein concentrations were determined by Bradford protein assay (Thermo Fisher Scientific). Equal amounts of lysates were loaded for individual experiments and β-tubulin was immunoblotted as the loading control. HRP chemiluminescence images were captured with the ChemiDoc Touch Imaging System (Bio-Rad). For quantification, 3 to 5 independent repeats were performed, and data were analyzed using the Image Lab software.

### Immunohistochemistry

Postnatal mouse brains were perfusion-fixed in 4% paraformaldehyde (PFA) at 4°C overnight. Tissues were dehydrated in 30% sucrose solution for 36-48 h and embedded in the Tissue-Tek O.C.T compound (Sakura). Coronal sections (30-μm thick) were prepared by cryostat sectioning. Brain sections were completely dried at RT, washed with PBS and antigen-retrieved in 10 mM citrate acid (pH 6.0) at 95°C for 20 min. The sections were brought to RT, washed with PBS for 3 times and permeabilized in blocking solution (0.1% Triton X-100, 4% normal goat serum or normal donkey serum in PBS) for 1 h at RT. Samples were then incubated with primary antibodies at 4°C overnight, followed by fluorescent secondary antibodies for 2 h at RT. Images were acquired using the Olympus VS120 virtual slide system (20 X tile scan) or the ANDOR spinning disc confocal microscope with a 60 X objective.

The myelinated area in outlined corpus callosum was measured by the MBP signals above an empirically determined threshold applied to all samples. The length of myelin sheath in the S1 cortex was averaged from MBP-labeled axons in 12 to 24 squares (500 × 500 pixels) from 6 to 12 brain sections per animal. The densities of total OLs (OLIG2^+^), OPCs (OLIG2^+^ PDGFRA^+^) and mature OLs (OLIG2^+^ CC1^+^) were averaged from 8 to 12 brain sections per animal. At least three pairs of littermates were analyzed for each measurement, and the brain sections used for quantification covered the rostral and ventral tips of the corpus callosum. All immunohistochemistry images were analyzed with ImageJ and MetaMorph.

### RNAscope and microRNAscope

Cryo-sections of postnatal brain tissues were prepared as above. The RNAScope^®^ Multiplex Fluorescent Reagent v2 kit and the miRNAscope™ HD Reagent Kit (ACDBio) were used to label mRNAs and miRNAs, respectively, following the manufacturer’s manuals. Co-labelling of OLIG2 with MiR7a-5p was achieved using the RNA-Protein Co-detection Kit (ACDBio). All probes were ordered from ACDBio and detailed information is shown in Table S4.

Nuclei were stained with DAPI (Sigma-Aldrich). Images were acquired with the Olympus VS120 virtual slide system (20 X tile scan) or the Olympus spinning disc confocal microscope (20 X tile scan). ImageJ and MetaMorph were used for image analyses. For quantification of mRNA levels, an oval of fixed size encompassing DAPI and all other fluorescent signals was drawn for each cell, and the integrated fluorescence intensity within the oval was determined for 40 cells in the corpus callosum.

### Transmission electron microscopy (TEM)

P14 brains were fixed in 4% PFA and post-fixed in 2.5% glutaraldehyde (in 0.1 M phosphate buffer, pH 7.4) at 4°C overnight. One-millimeter-thick slices were cut with a brain slicer matrix and razor blades. The samples were washed in phosphate buffer (pH 7.4) and dehydrated in increasing concentrations of ethanol (30%, 50%, 70%, 90%, and 100% for 3 times). After embedding in Epon-filled molds, the samples were sectioned at 85 nm of thickness by an ultramicrotome (Leica, UC7) and viewed under a Spirit 120 kV transmission electron microscope (Thermo Fisher Scientific). The thickness of myelin sheath was measured by MetaMorph.

### iBAQ quantitative mass spectrometry and GO analyses

Forebrains were dissected from *Z^f/f^* and *Z^f/f^;Nestin-Cre* littermates at P0 and P14 and snap-frozen in liquid nitrogen immediately. Tissue lysates were prepared and processed as previously described (*76*). Liquid chromatography-mass spectrometry (LC-MS) analyses of trypsin-digested peptides were performed on a Q Exactive HF-X hybrid Quadrupole-Orbitrap instrument (Thermo Fisher Scientific) coupled with an Easy-nLC 1200 system. MS/MS spectra were extracted by resolving from RAW files. Collected data were analyzed using the iBAQ (intensity-based absolute quantification) method as implemented in MaxQuant software (https://maxquant.org/maxquant/, version 2.1.4.0). RAW file datasets were searched against the mouse proteome downloaded from UniProt. Search parameters were: 5% false discovery rate (FDR) at the peptide-spectrum match (PSM) level, 10 ppm precursor mass tolerance, 20 ppm fragment mass tolerance, variable modification Cys 57.02146, and three maximum number of missed cleavage sites. Up- and down-regulated proteins were identified using adjusted iBAQ intensity ratios defined as follows:

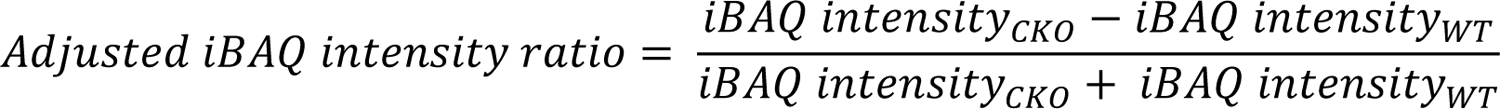

A protein with an adjusted iBAQ intensity ratio ≥ 0.15 or ≤ −0.15 was considered up- or down-regulated. Only proteins that showed the same direction of change in both absolute intensity and iBAQ intensity across all CKO-WT pairs were considered “correctly detected” and retained for further analysis.

Gene Ontology (GO) enrichment analysis was performed using the R package clusterProfiler (version 4.6.0). The p value was adjusted by the Benjamini–Hochberg method. Enriched GO terms with false discovery rate (FDR) < 0.05 were selected. Visualization of GO results were achieved by the R package (version 4.2.2). Network analysis of up- and down-regulated proteins in P14 mouse brains was performed by igraph R package (version 1.4.1). Venn diagrams were generated using the jvenn plug-in (http://jvenn.toulouse.inra.fr/app/index.html).

### Small RNA sequencing (sRNA-Seq) and analysis

Whole brains were dissected from *Z^f/f^* and *Z^f/f^;Nestin-Cre* littermates at P2 and frozen in liquid nitrogen immediately. Sample processing and sRNA-Seq was done by BGI Genomics (Shenzhen, China). DNBSEQ was used for sequencing and clean data were aligned to the mouse reference genome (GRCm38/mm10) using HISAT and to sRNAs in miRbase (https://www.mirbase.org). miRDeep2 was used to predicted newly discovered miRNAs. miRNA expression was quantified with Unique Molecular Identifiers (UMI), and DEGseq was used for analysis of differentially expressed genes defined as | log_2_(fold-change) | > 0 and P_adj_ < 0.05 (Benjamini-Hochberg).

### Computational analysis of intrinsically disordered regions

IDR prediction was performed by selected algorithms from the D^2^P^2^ database (https://d2p2.pro/) (*43*). For each protein, the total number of amino acids within IDRs was calculated by C# and divided by the protein length to obtain % of IDR. Then these proteins are categorized into four groups based the results (IDR % < 5%, 5-50%, 50-95% and ≥ 95%). The number of large IDRs (> 30 consecutive amino acids) in each protein was also counted, and the fraction of proteins with a certain number of IDRs was fitted with a fifth-order polynomial curve (R^2^ >0. 99) using GraphPad Prism 9.

### Stereoscopic injection

An anesthetized P3 pup was immobilized on a stereotaxic device (RWD Life Science) with its head position fixed by the ear rods on both sides. The Lambda of the posterior fontanel served as the origin of the three-dimensional coordinate system of the skull. The syringe needle was slowly and vertically moved to the target site (bilateral ventricular coordinates are X = ± 0.8 mm, Y = 0.8 mm, Z = −1.9 mm) and inserted into the cerebral ventricle. Bilateral injections of agomirs (0.05 nmol in 500 nL) or antagomirs (0.1 nmol in 500 nL) were performed at 50 nL/sec into the subventricular zone region for each animal. After injection, the needle was left in position for 5 min before being slowly removed. Then these pups were labeled by toe-clipping (tissues saved for genotyping) and kept with the dam until they were ready for brain collection on P10 (agomirs) or P14 (antagomirs).

### Data analysis and statistics

Data are shown as mean ± SEM unless otherwise noted. Comparison of two groups was performed by two-tailed Student’s unpaired *t*-test and multiple comparisons were performed by one-way ANOVA using Graph Prism 6.0.

## Acknowledgement

We thank Dr. Mengsheng Qiu (Hangzhou Normal University), Dr. Zhihua Gao (Zhejiang University) and Dr. Chong Liu (Zhejiang University) for providing reagents and valuable suggestions. We thank Dr. Cheng Ma and Dr. Liyan Wang from the Protein facility at CFZSM (Core facilities, Zhejiang University School of Medicine) for technical assistance. We are grateful for microscopy support from Dr. Junli Xuan and Dr. Sanhua Fang at CFZSM. Animal care for this study was provided by the Laboratory Animal Center at Zhejiang University.

## Author contributions

Conceptualization and supervision: X.G. and Z.W.

Experimental execution and data analysis: J.L., S.Z., R.F., X.S., G.W., J.G., S.X. and L.Z.

Resources: X.G., Z.W., S.D., B.Y., J.J. and A.R.

Visualization: J.L. and S.M.

Writing – original draft: J.L., X.S., X.G. and Z.W.

Writing – review & editing: J.L., S.Z., X.G. and Z.W.

---

**Figure S1.**
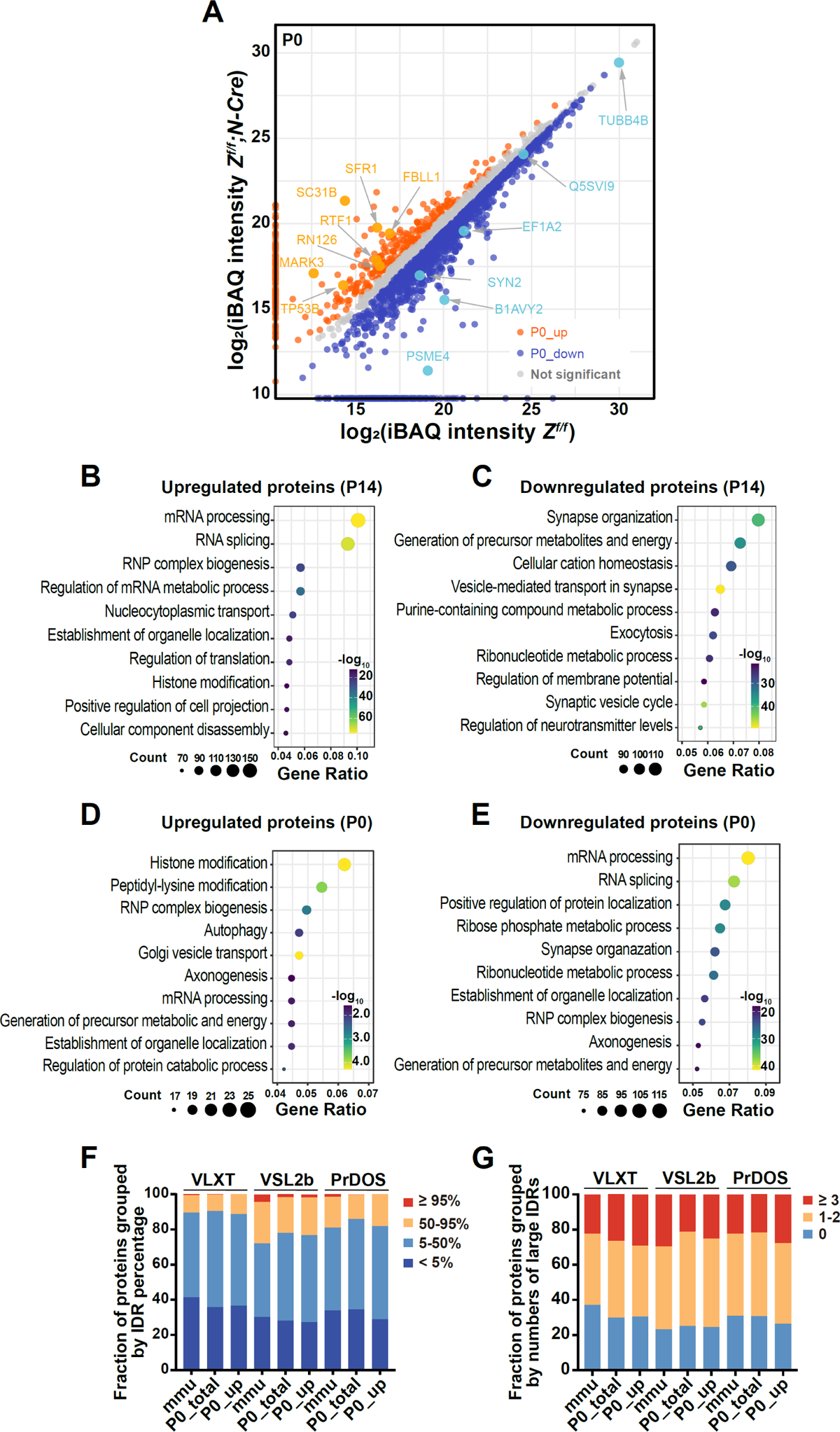
Analyses of proteomic changes in *Z^f/f^;N-Cre* forebrains at P0 and P14. **(A)** P0 forebrain homogenates from *Z^f/f^* and *Z^f/f^;N-Cre* animals (N = 3 pairs of littermates) were analyzed by mass spectrometry. Representatives of differentially expressed proteins in *Z^f/f^;N-Cre* mice are indicated by arrows. **(B, C)** The top 10 most enriched GO terms (biological processes) with regard to upregulated proteins (B) or downregulated proteins (C) in *Z^f/f^;N-Cre* forebrain at P14. **(D, E)** The top 10 most enriched GO terms (biological processes) with regard to upregulated proteins (D) or downregulated proteins (E) in *Z^f/f^;N-Cre* forebrain at P0. **(F)** Proteins in the total mouse proteome (*mmu*), in the forebrain proteome from control mice at P0 (P0_total), and those upregulated in *Z^f/f^;N-Cre* forebrain samples at P0 (P0_up) were ranked based on their IDR contents calculated by three algorithms (VXLT, VSL2b and PrDOS). **(G)** As in (F), proteins were ranked based on the number of large IDRs they harbor.

**Figure S2.**
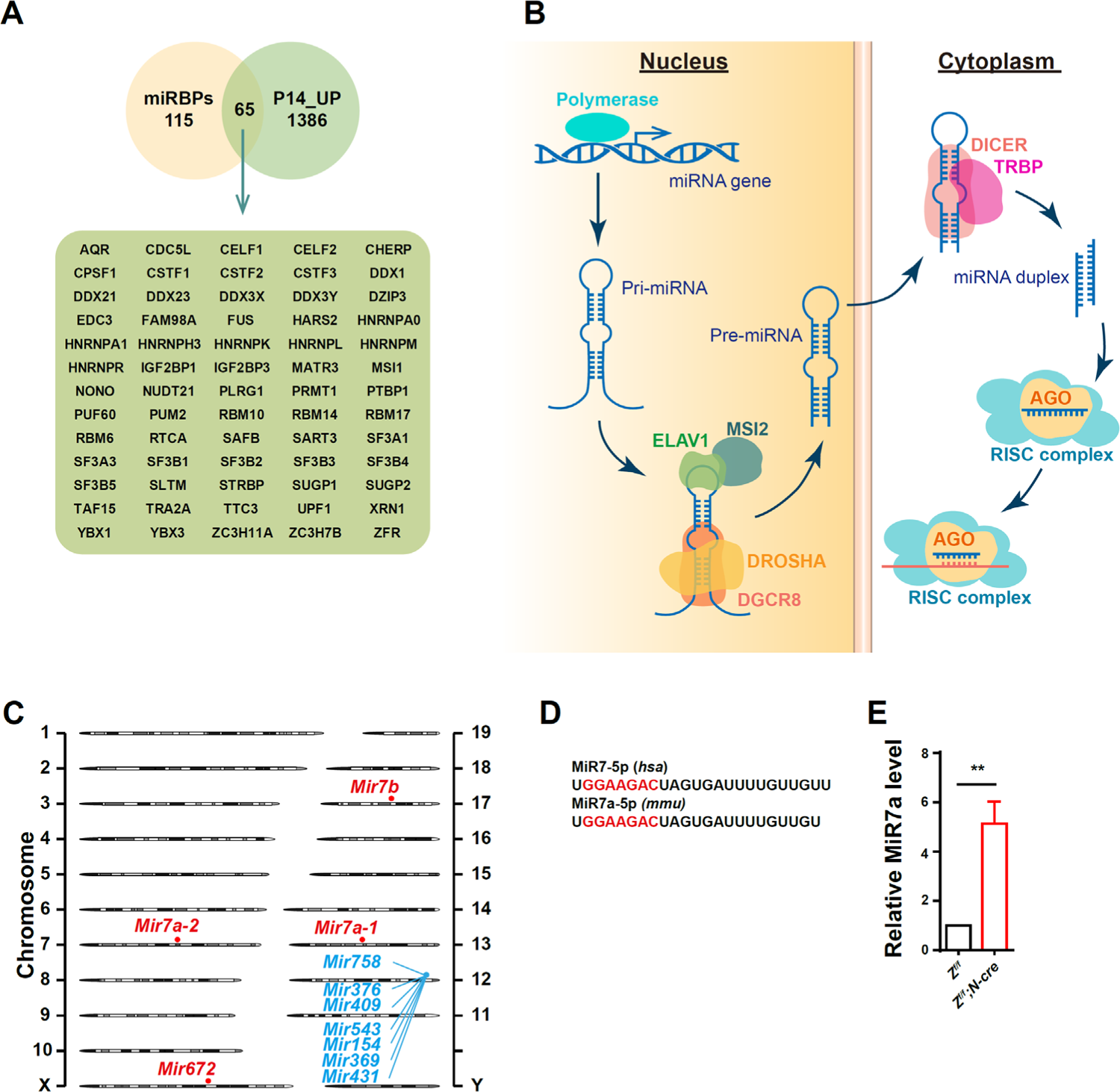
Deletion of ZSWIM8 causes miRBP accumulation and MiR7 upregulation. **(A)** Venn diagram showing one third of experimentally validated miRBPs from a previous study (*1*) among upregulated proteins in the P14 *Z^f/f^;N-Cre* brain (P14_UP). **(B)** A schematic of the miRNA biogenesis pathway (created with BioRender, https://www.biorender.com). ELAV1 and MSI2 are known to regulate MiR7 processing. Pri-miRNA, primary miRNA; pre-miRNA, precursor miRNA. **(C)** Chromosome mapping of the 10 miRNAs upregulated in *Z^f/f^;N-Cre* mice. **(D)** The seed sequence of MiR7 (shown in red) is conserved between human and mouse. **(E)** Validation of MiR7 upregulation in *Z^f/f^;N-Cre* whole brain by real-time qRT-PCR. Samples were collected from 3 pairs of littermates at P2. **p < 0.01 (Student’s unpaired *t*-test).

**Figure S3.**
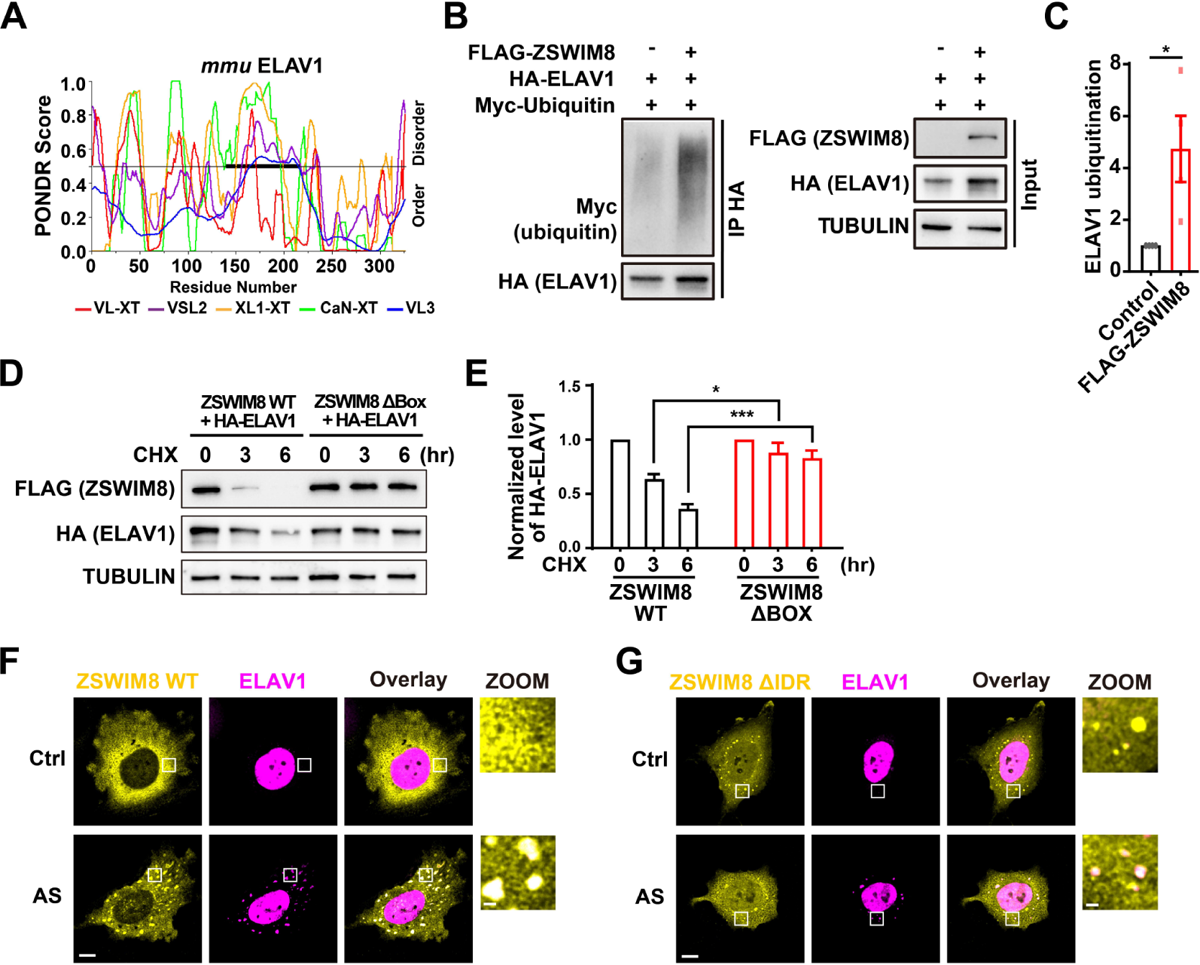
ZSWIM8 promotes ELAV1 degradation by a “disorder-targets-misorder” mechanism. **(A)** IDR prediction results of mouse ELAV1 from PONDR. The thick black line indicates a long IDR detected by all five algorithms. **(B)** HEK293FT cells were transfected with the indicated constructs and treated with bortezomib (5 μM, 4h). HA-ELAV1 was immunoprecipitated by anti-HA nanobody beads. Polyubiquitination signals were detected by anti-MYC antibodies. **(C)** Four independent experiments as in (B) were used for quantification of ELAV1 ubiquitination. *p < 0.05 (Student’s unpaired *t*-test). **(D)** HEK293FT cells were transfected with the indicated constructs and treated with 50 μg/ml cycloheximide (CHX) for 0, 3 and 6 h before harvest. Equal amounts of lysates were blotted with the indicated antibodies. **(E)** Four independent experiments as in (D) were used for quantification of ELAV1 levels. *p < 0.05 and ***p < 0.001 (Student’s unpaired *t*-test). **(F, G)** Confocal images of HUVEC cells co-transfected with HA-tagged ELAV1 and V5-tagged ZSWIM8-WT (F) or ZSWIM8-ΔIDR (G). Scale bar, 10 μm. Sodium arsenite (500 μM, 30 min) was used to induce oxidative stress. Zoom-in images of the boxed regions are shown on the right. Scale bar, 2 μm.

**Figure S4.**
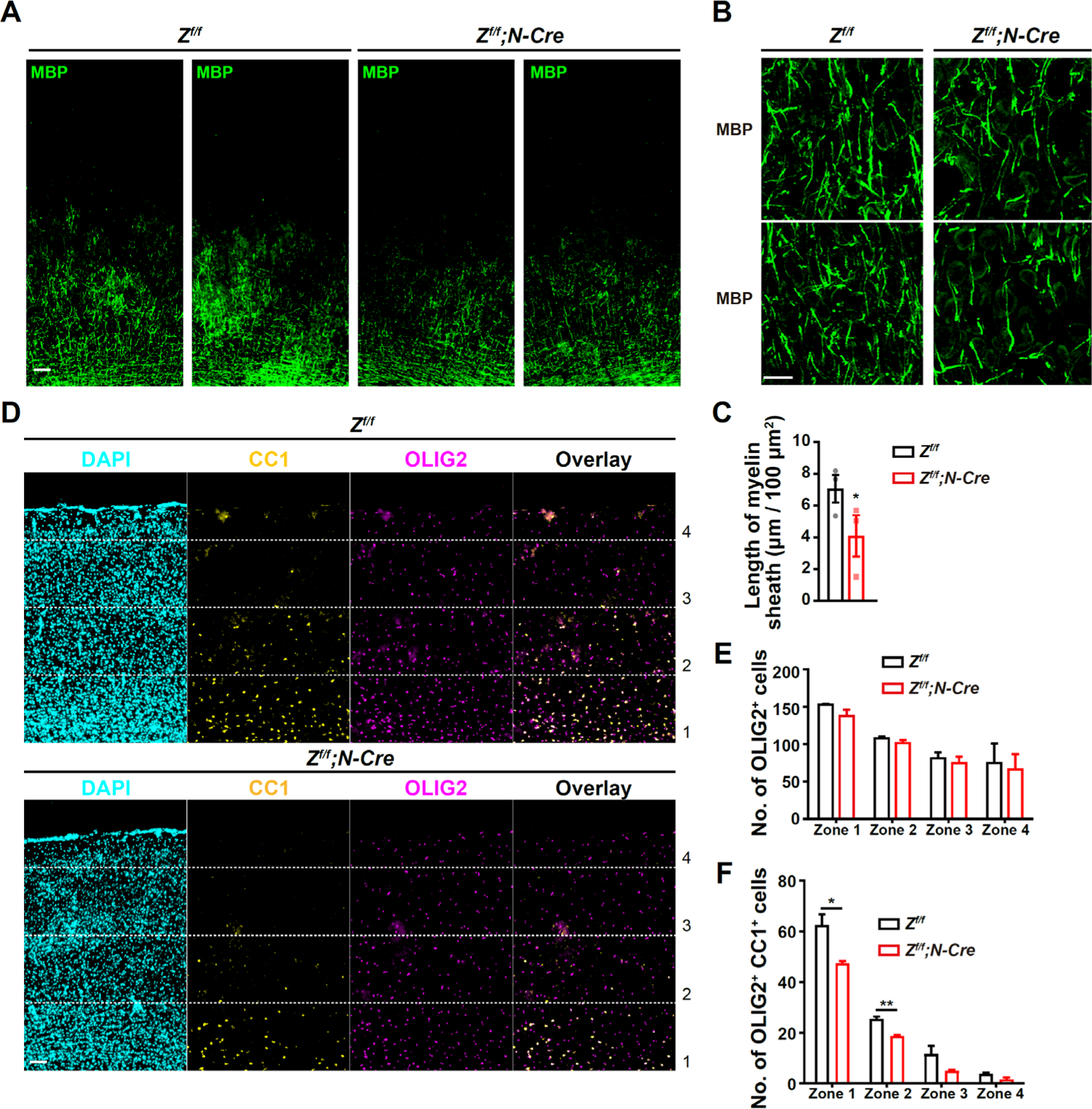
Myelination defects in S1 cortex of *Z^f/f^;N-Cre* mice. (**A**) Immunohistochemistry of MBP in the S1 cortex (coronal sections) from *Z^f/f^* and *Z^f/f^;N-Cre* animals at P14. Scale bar, 20 μm. **(B, C)** High magnification (60 X) images of anti-MBP IHC in layers IV-V of the S1 cortex) from *Z^f/f^* and *Z^f/f^;N-Cre* animals at P14. Scale bar, 20 μm. Quantification of the myelin sheath density in these regions was shown in (C). N = 3 pairs of littermates. *p < 0.05 (Student’s unpaired *t*-test). **(D)** Anti-OLIG2 and anti-CC1 IHC of the S1 cortex of *Z^f/f^* and *Z^f/f^;N-Cre* animals at P14. DAPI labels all nuclei. Scale bar, 20 μm. The imaged areas were equally divided into four zones for the ease of quantification, with zone 1 being the deepest layer of the cortex. **(E, F)** Numbers of total OL lineage cells (OLIG2^+^, E) and mature OLs (OLIG2^+^ CC1^+^, F) were quantified for each zone of the S1 cortex based on IHC results from (D). N = 3 pairs of littermates. *p < 0.05 and **p < 0.01 relative to indicated controls (Student’s unpaired *t*-test)

**Figure S5.**
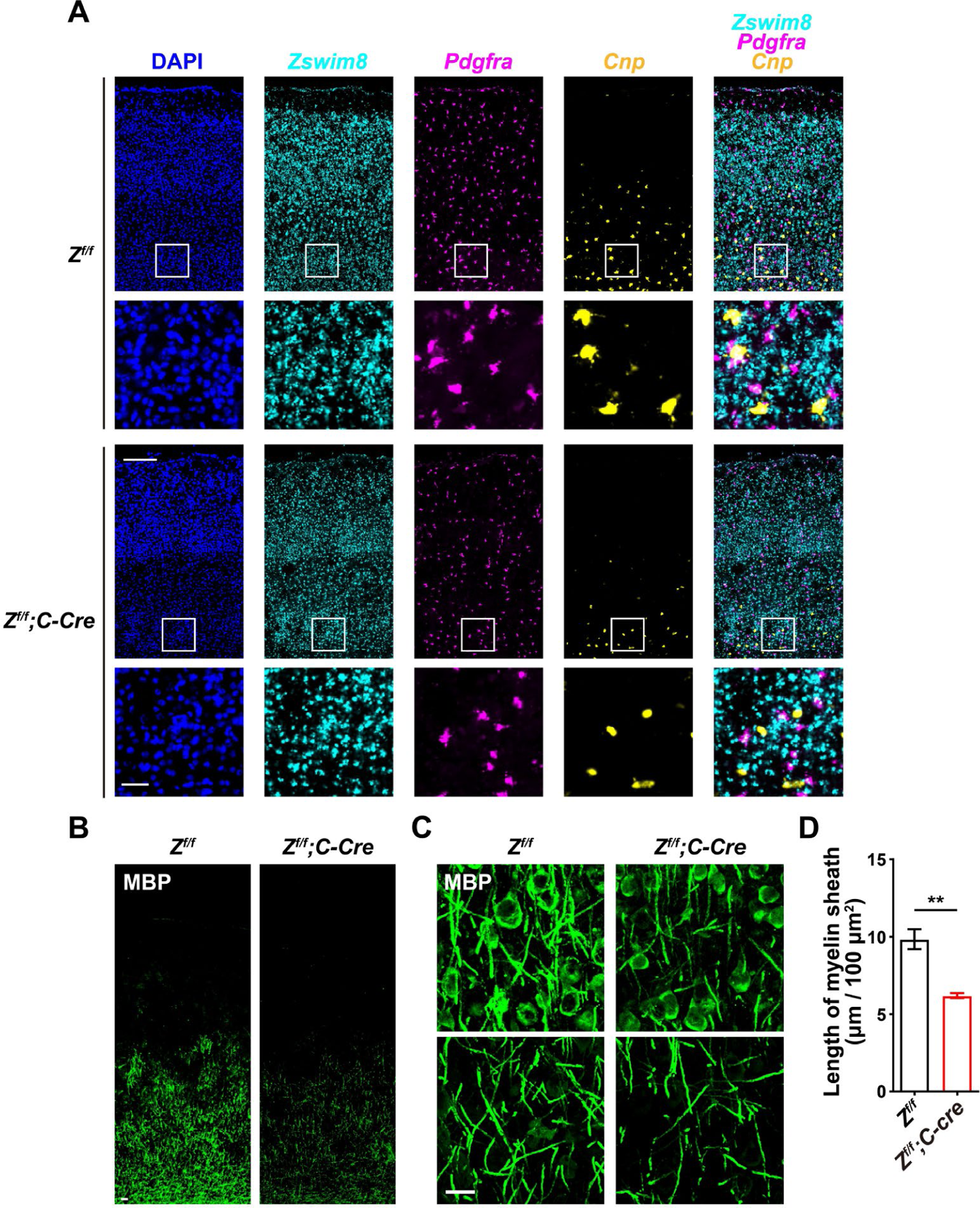
Conditional deletion of ZSWIM8 in premyelinating OLs impairs OL maturation and myelination in the S1 cortex. **(A)** Representative RNAscope images of *Zswim8*, *Pdgfra* and *Cnp* in the S1 cortex of P14 *Z^f/f^* and *Z^f/f^;C-Cre* mice. Scale bar, 50 μm. Zoom-in images of the boxed regions are shown below (Scale bar = 10 μm). **(B)** Representative images of MBP immunohistochemistry in the S1 cortex (coronal sections) from *Z^f/f^* and *Z^f/f^;C-Cre* animals at P14. Scale bar, 10 μm. **(C)** High magnification (60 X) images of MBP immunohistochemistry in layers IV-V of the S1 corte from *Z^f/f^* and *Z^f/f^;C-Cre* animals at P14. Scale bar, 20 μm. **(D)** Quantification of the myelin sheath density in S1 cortex layers IV-V. N = 3 pairs of littermates. **p < 0.01 (Student’s unpaired *t*-test).

**Figure S6.**
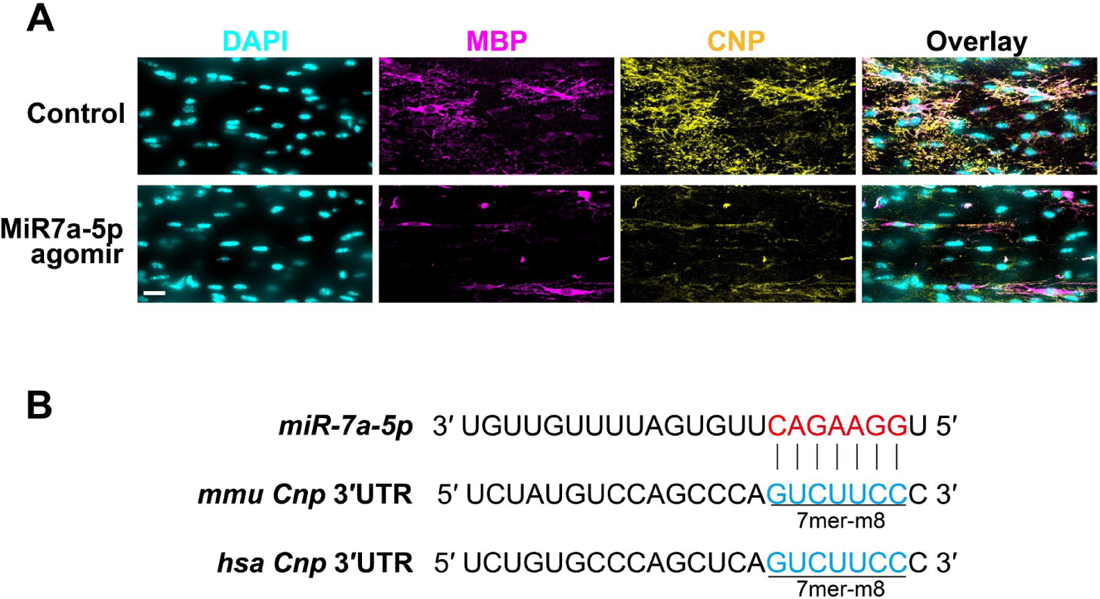
Excessive MiR7 inhibits myelination in the corpus callosum. **(A)** Representative MBP immunohistochemistry images of the corpus callosum from mice injected with control or MiR7 agomir. Samples were collected at P14. Nuclei were stained by DAPI. Scale bar, 20 μm. **(B)** Human and mouse Cnp 3’UTRs contain a conserved *MiR7a-5p* seed sequence.

**Figure S7.**
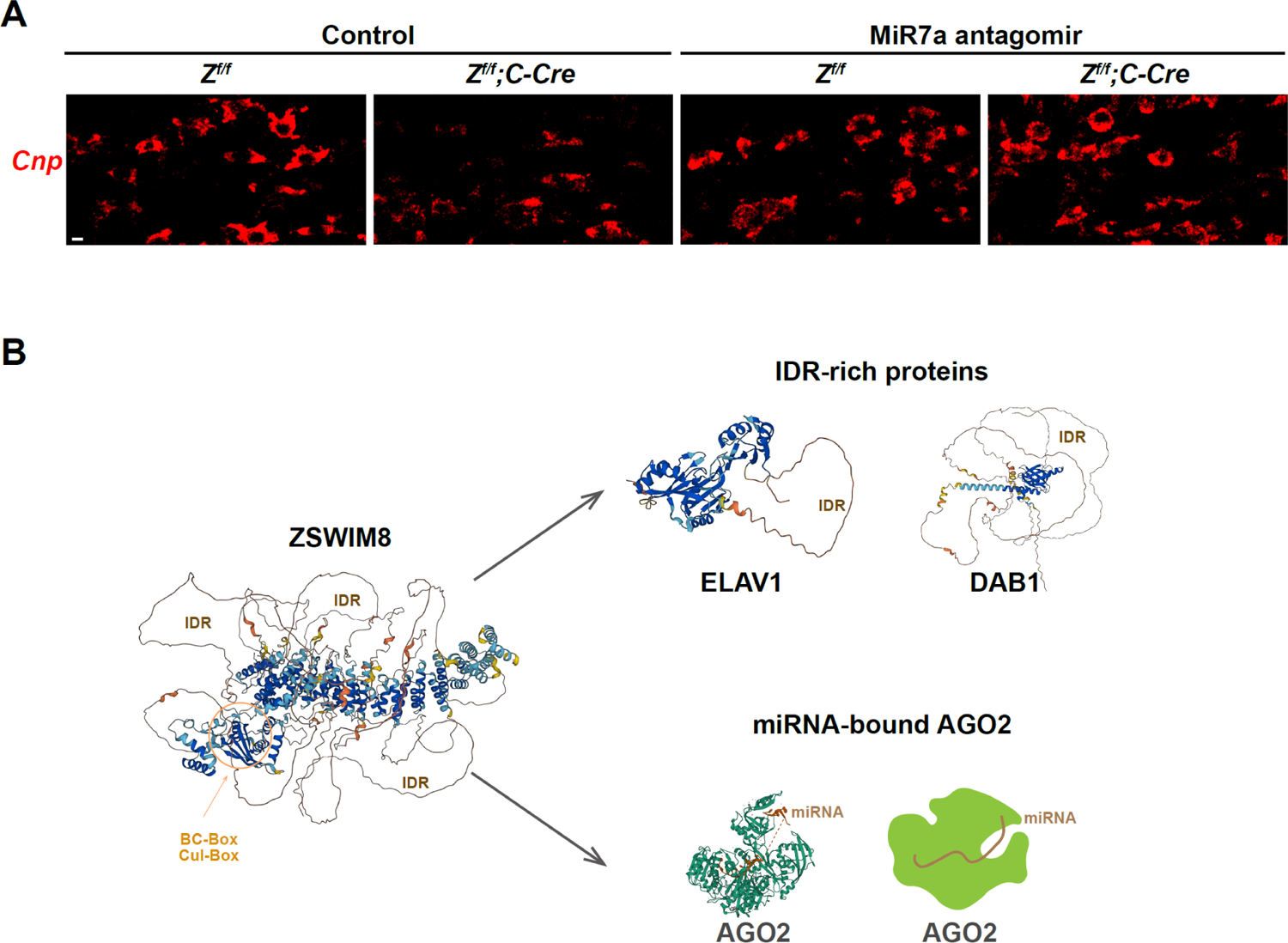
MiR7 functions downstream of ZSWIM8 and facilitates AGO2 degradation. **(A)** *Cnp* mRNA detected by RNAscope in the corpus callosum of P14 *Z^f/f^* and *Z^f/f^;C-Cre* animals injected with control or MiR7a antagomir. Representative images selected from 3 pairs of littermates. Scale bar, 10 μm. **(B)** ZSWIM8 binds typical IDR-containing proteins such as ELAV1 and DAB1 using the “disorder-targets-misorder” mechanism. On the other hand, ZSWIM8 can also recognize AGO2 in a miRNA-dependent manner. Predicted structures of ZSWIM8, ELAV1 and DAB1 were generated by AlphaFold (*2, 3*). The structure of MIR20a-bound hAGO2 is adapted from Protein Data Bank (PDB: 4F3T).

## Reference

1. B. Tsang, I. Pritisanac, S. W. Scherer, A. M. Moses, J. D. Forman-Kay, Phase Separation as a Missing Mechanism for Interpretation of Disease Mutations. Cell 183, 1742–1756 (2020).

2. J. J. Ward, J. S. Sodhi, L. J. McGuffin, B. F. Buxton, D. T. Jones, Prediction and functional analysis of native disorder in proteins from the three kingdoms of life. J Mol Biol 337, 635–645 (2004).

3. S. Vucetic, C. J. Brown, A. K. Dunker, Z. Obradovic, Flavors of protein disorder. Proteins 52, 573–584 (2003).

4. C. J. Oldfield, A. K. Dunker, Intrinsically disordered proteins and intrinsically disordered protein regions. Annu Rev Biochem 83, 553–584 (2014).

5. W. Basile, M. Salvatore, C. Bassot, A. Elofsson, Why do eukaryotic proteins contain more intrinsically disordered regions? PLoS Comput Biol 15, e1007186 (2019).

6. B. Shen et al., Computational Screening of Phase-separating Proteins. Genomics Proteomics Bioinformatics 19, 13–24 (2021).

7. L. D. Gallego et al., Phase separation directs ubiquitination of gene-body nucleosomes. Nature 579, 592–597 (2020).

8. A. Zbinden, M. Perez-Berlanga, P. De Rossi, M. Polymenidou, Phase Separation and Neurodegenerative Diseases: A Disturbance in the Force. Dev Cell 55, 45–68 (2020).

9. A. Castello et al., Insights into RNA biology from an atlas of mammalian mRNA-binding proteins. Cell 149, 1393–1406 (2012).

10. A. Castello et al., Comprehensive Identification of RNA-Binding Domains in Human Cells. Mol Cell 63, 696–710 (2016).

11. M. W. Hentze, A. Castello, T. Schwarzl, T. Preiss, A brave new world of RNA-binding proteins. Nat Rev Mol Cell Biol 19, 327–341 (2018).

12. B. M. Beckmann et al., The RNA-binding proteomes from yeast to man harbour conserved enigmRBPs. Nat Commun 6, 10127 (2015).

13. P. H. Sudmant, H. Lee, D. Dominguez, M. Heiman, C. B. Burge, Widespread Accumulation of Ribosome-Associated Isolated 3’ UTRs in Neuronal Cell Populations of the Aging Brain. Cell Rep 25, 2447–2456 e2444 (2018).

14. O. Mauger, F. Lemoine, P. Scheiffele, Targeted Intron Retention and Excision for Rapid Gene Regulation in Response to Neuronal Activity. Neuron 92, 1266–1278 (2016).

15. A. Molliex et al., Phase separation by low complexity domains promotes stress granule assembly and drives pathological fibrillization. Cell 163, 123–133 (2015).

16. P. Connerty, A. Ahadi, G. Hutvagner, RNA Binding Proteins in the miRNA Pathway. Int J Mol Sci 17, (2015).

17. J. Trendel et al., The Human RNA-Binding Proteome and Its Dynamics during Translational Arrest. Cell 176, 391–403 e319 (2019).

18. S. Maharana et al., RNA buffers the phase separation behavior of prion-like RNA binding proteins. Science 360, 918–921 (2018).

19. A. Patel et al., A Liquid-to-Solid Phase Transition of the ALS Protein FUS Accelerated by Disease Mutation. Cell 162, 1066–1077 (2015).

20. J. A. Kolhe, N. L. Babu, B. C. Freeman, The Hsp90 molecular chaperone governs client proteins by targeting intrinsically disordered regions. Mol Cell 83, 2035–2044 e2037 (2023).

21. C. Pohl, I. Dikic, Cellular quality control by the ubiquitin-proteasome system and autophagy. Science 366, 818–822 (2019).

22. R. T. Timms et al., Defining E3 ligase-substrate relationships through multiplex CRISPR screening. Nat Cell Biol, (2023).

23. Z. Zhang et al., Elucidation of E3 ubiquitin ligase specificity through proteome-wide internal degron mapping. Mol Cell 83, 3377–3392 e3376 (2023).

24. Z. Wang et al., The EBAX-type Cullin-RING E3 ligase and Hsp90 guard the protein quality of the SAX-3/Robo receptor in developing neurons. Neuron 79, 903–916 (2013).

25. J. C. Rosenbaum et al., Disorder targets misorder in nuclear quality control degradation: a disordered ubiquitin ligase directly recognizes its misfolded substrates. Mol Cell 41, 93–106 (2011).

26. D. C. Stieg et al., A complex molecular switch directs stress-induced cyclin C nuclear release through SCF(Grr1)-mediated degradation of Med13. Mol Biol Cell 29, 363–375 (2018).

27. V. Narayan, E. Pion, V. Landre, P. Muller, K. L. Ball, Docking-dependent ubiquitination of the interferon regulatory factor-1 tumor suppressor protein by the ubiquitin ligase CHIP. J Biol Chem 286, 607–619 (2011).

28. V. Vamadevan, N. Chaudhary, S. Maddika, Ubiquitin-assisted phase separation of dishevelled-2 promotes Wnt signalling. J Cell Sci 135, (2022).

29. G. Wang et al., The ZSWIM8 ubiquitin ligase regulates neurodevelopment by guarding the protein quality of intrinsically disordered Dab1. Cereb Cortex 33, 3866–3881 (2023).

30. E. R. Kingston, L. W. Blodgett, D. P. Bartel, Endogenous transcripts direct microRNA degradation in Drosophila, and this targeted degradation is required for proper embryonic development. Mol Cell 82, 3872–3884 e3879 (2022).

31. C. Y. Shi et al., ZSWIM8 destabilizes many murine microRNAs and is required for proper embryonic growth and development. Genome Res, (2023).

32. B. T. Jones et al., Target-directed microRNA degradation regulates developmental microRNA expression and embryonic growth in mammals. Genes Dev, (2023).

33. C. Y. Shi et al., The ZSWIM8 ubiquitin ligase mediates target-directed microRNA degradation. Science 370, (2020).

34. J. Han et al., A ubiquitin ligase mediates target-directed microRNA decay independently of tailing and trimming. Science 370, (2020).

35. J. Liu et al., Argonaute2 is the catalytic engine of mammalian RNAi. Science 305, 1437–1441 (2004).

36. G. Hutvagner, M. J. Simard, Argonaute proteins: key players in RNA silencing. Nat Rev Mol Cell Biol 9, 22–32 (2008).

37. G. Meister, Argonaute proteins: functional insights and emerging roles. Nat Rev Genet 14, 447–459 (2013).

38. J. Zhao et al., MicroRNA-7: expression and function in brain physiological and pathological processes. Cell Biosci 10, 77 (2020).

39. J. L. Horsham et al., MicroRNA-7: A miRNA with expanding roles in development and disease. Int J Biochem Cell Biol 69, 215–224 (2015).

40. A. Pollock, S. Bian, C. Zhang, Z. Chen, T. Sun, Growth of the developing cerebral cortex is controlled by microRNA-7 through the p53 pathway. Cell Rep 7, 1184–1196 (2014).

41. L. Zhang et al., Counter-Balance Between Gli3 and miR-7 Is Required for Proper Morphogenesis and Size Control of the Mouse Brain. Front Cell Neurosci 12, 259 (2018).

42. X. Zhao, J. Wu, M. Zheng, F. Gao, G. Ju, Specification and maintenance of oligodendrocyte precursor cells from neural progenitor cells: involvement of microRNA-7a. Mol Biol Cell 23, 2867–2878 (2012).

43. M. E. Oates et al., D(2)P(2): database of disordered protein predictions. Nucleic Acids Res 41, D508–516 (2013).

44. F. Cunningham et al., Ensembl 2022. Nucleic Acids Res 50, D988–D995 (2022).

45. T. Zarin et al., Proteome-wide signatures of function in highly diverged intrinsically disordered regions. Elife 8, (2019).

46. R. van der Lee et al., Classification of intrinsically disordered regions and proteins. Chem Rev 114, 6589–6631 (2014).

47. T. Treiber et al., A Compendium of RNA-Binding Proteins that Regulate MicroRNA Biogenesis. Mol Cell 66, 270–284 e213 (2017).

48. N. R. Choudhury et al., Tissue-specific control of brain-enriched miR-7 biogenesis. Genes Dev 27, 24–38 (2013).

49. X. C. Fan, J. A. Steitz, Overexpression of HuR, a nuclear-cytoplasmic shuttling protein, increases the in vivo stability of ARE-containing mRNAs. Embo j 17, 3448–3460 (1998).

50. K. Arimoto, H. Fukuda, S. Imajoh-Ohmi, H. Saito, M. Takekawa, Formation of stress granules inhibits apoptosis by suppressing stress-responsive MAPK pathways. Nat Cell Biol 10, 1324–1332 (2008).

51. S. Xu et al., Cytosolic stress granules relieve the ubiquitin-proteasome system in the nuclear compartment. EMBO J 42, e111802 (2023).

52. N. T. Schirle, I. J. MacRae, The crystal structure of human Argonaute2. Science 336, 1037–1040 (2012).

53. E. Elkayam et al., The structure of human argonaute-2 in complex with miR-20a. Cell 150, 100–110 (2012).

54. J. Sheu-Gruttadauria et al., Structural Basis for Target-Directed MicroRNA Degradation. Mol Cell 75, 1243–1255 e1247 (2019).

55. M. Quevillon Huberdeau et al., Phosphorylation of Argonaute proteins affects mRNA binding and is essential for microRNA-guided gene silencing in vivo. EMBO J 36, 2088–2106 (2017).

56. N. T. Schirle, J. Sheu-Gruttadauria, I. J. MacRae, Structural basis for microRNA targeting. Science 346, 608–613 (2014).

57. R. J. Golden et al., An Argonaute phosphorylation cycle promotes microRNA-mediated silencing. Nature 542, 197–202 (2017).

58. B. Elbaz, B. Popko, Molecular Control of Oligodendrocyte Development. Trends Neurosci 42, 263–277 (2019).

59. C. Hayashi, N. Suzuki, Heterogeneity of Oligodendrocytes and Their Precursor Cells. Adv Exp Med Biol 1190, 53–62 (2019).

60. W. D. Richardson, N. Kessaris, N. Pringle, Oligodendrocyte wars. Nat Rev Neurosci 7, 11–18 (2006).

61. G. La Manno et al., Molecular architecture of the developing mouse brain. Nature 596, 92–96 (2021).

62. S. Kuhn, L. Gritti, D. Crooks, Y. Dombrowski, Oligodendrocytes in Development, Myelin Generation and Beyond. Cells 8, (2019).

63. P. Sheng et al., Screening of Drosophila microRNA-degradation sequences reveals Argonaute1 mRNA’s role in regulating miR-999. Nat Commun 14, 2108 (2023).

64. L. Li et al., Widespread microRNA degradation elements in target mRNAs can assist the encoded proteins. Genes Dev 35, 1595–1609 (2021).

65. E. R. Kingston, D. P. Bartel, Ago2 protects Drosophila siRNAs and microRNAs from target-directed degradation, even in the absence of 2’-O-methylation. Rna 27, 710–724 (2021).

66. B. F. Donnelly et al., The developmentally timed decay of an essential microRNA family is seed-sequence dependent. Cell Rep 40, 111154 (2022).

67. S. Kumar, A. Downie Ruiz Velasco, G. Michlewski, Oleic Acid Induces MiR-7 Processing through Remodeling of Pri-MiR-7/Protein Complex. J Mol Biol 429, 1638–1649 (2017).

68. Y. C. Lu et al., ELAVL1 modulates transcriptome-wide miRNA binding in murine macrophages. Cell Rep 9, 2330–2343 (2014).

69. S. Huang et al., Loss of Super-Enhancer-Regulated circRNA Nfix Induces Cardiac Regeneration After Myocardial Infarction in Adult Mice. Circulation 139, 2857–2876 (2019).

70. L. Shi et al., A tumor-suppressive circular RNA mediates uncanonical integrin degradation by the proteasome in liver cancer. Sci Adv 7, (2021).

71. J. H. Yoon et al., Scaffold function of long non-coding RNA HOTAIR in protein ubiquitination. Nat Commun 4, 2939 (2013).

72. B. Kleaveland, C. Y. Shi, J. Stefano, D. P. Bartel, A Network of Noncoding Regulatory RNAs Acts in the Mammalian Brain. Cell 174, 350–362 e317 (2018).

73. F. Tronche et al., Disruption of the glucocorticoid receptor gene in the nervous system results in reduced anxiety. Nat Genet 23, 99–103 (1999).

74. C. Lappe-Siefke et al., Disruption of Cnp1 uncouples oligodendroglial functions in axonal support and myelination. Nat Genet 33, 366–374 (2003).

75. G. L. Wieser et al., Neuroinflammation in white matter tracts of Cnp1 mutant mice amplified by a minor brain injury. Glia 61, 869–880 (2013).

76. X. Kong et al., Fine-tuning of mTOR signaling by the UBE4B-KLHL22 E3 ubiquitin ligase cascade in brain development. Development 149, (2022).

## References

1. T. Treiber et al., A Compendium of RNA-Binding Proteins that Regulate MicroRNA Biogenesis. Mol Cell 66, 270–284 e213 (2017).

2. J. Jumper et al., Highly accurate protein structure prediction with AlphaFold. Nature 596, 583–589 (2021).

3. M. Varadi et al., AlphaFold Protein Structure Database: massively expanding the structural coverage of protein-sequence space with high-accuracy models. Nucleic Acids Res 50, D439–D444 (2022).

